# Distinct value computations support rapid sequential decisions

**DOI:** 10.1101/2023.03.14.532617

**Authors:** Andrew Mah, Shannon S. Schiereck, Veronica Bossio, Christine M. Constantinople

**Author notes:** Zuckerman Institute, Columbia University; New York, NY 10027.

## Abstract

The value of the environment determines animals’ motivational states and sets expectations for error-based learning^1–3^. How are values computed? Reinforcement learning systems can store or “cache” values of states or actions that are learned from experience, or they can compute values using a model of the environment to simulate possible futures^3^. These value computations have distinct trade-offs, and a central question is how neural systems decide which computations to use or whether/how to combine them^4–8^. Here we show that rats use distinct value computations for sequential decisions within single trials. We used high-throughput training to collect statistically powerful datasets from 291 rats performing a temporal wagering task with hidden reward states. Rats adjusted how quickly they initiated trials and how long they waited for rewards across states, balancing effort and time costs against expected rewards. Statistical modeling revealed that animals computed the value of the environment differently when initiating trials versus when deciding how long to wait for rewards, even though these decisions were only seconds apart. Moreover, value estimates interacted via a dynamic learning rate. Our results reveal how distinct value computations interact on rapid timescales, and demonstrate the power of using high-throughput training to understand rich, cognitive behaviors.

## Introduction

There are many ways to compute value. Reinforcement learning provides a powerful framework for describing how animals or agents learn the value of different states and actions from experience and use those value estimates to guide behavior^3^. The value of the environment, or how much reward it is expected to yield, is important for motivation and sets expectations for reinforcement learning^1–3^.

There are many reinforcement learning methods for computing value that differ in their implementation, computational demands, and flexibility^3, 6, 8, 9^. For instance, some algorithms use a model of the world to flexibly estimate the value of states or actions by mental simulation or planning. Other algorithms cache values from direct experience, without an explicit model of the environment. These different reinforcement learning methods, which remarkably are thought to be supported by distinct neural circuits^10, 11^, have trade-offs between flexibility and computational efficiency^6, 8, 9^. They also represent two ends of a continuum^4–8^. A central question in neuroscience and psychology is determining how values are computed in animals including humans^12^. Moreover, neurobiologically-inspired value computations will likely lead to advances in next generation artificial intelligence^13^.

However, it is difficult to determine the value computations that subjects use, especially over behaviorally relevant timescales of seconds. In standard two-alternative forced choice tasks, the behavioral read-out is a binary choice, and the underlying values driving choice are obscure. State-of-the-art methods for revealing how values are computed use regression models that pool data over entire behavioral sessions^14^, or pre-determined subsets of trials^15^, thereby obscuring moment-by-moment changes in value computations. Therefore, whether or how multiple value computations interact on rapid timescales in the same subject is unclear.

## Results

### Rats’ deliberative and motivational decisions are sensitive to the value of the environment

We developed a temporal wagering task for rats, in which they were offered one of several water rewards on each trial, the volume of which (5, 10, 20, 40, 80µL) was indicated by a tone (Fig. 1a). The reward was assigned randomly to one of two ports, indicated by an LED. The rat could wait for an unpredictable delay to obtain the reward, or at any time could terminate the trial by poking in the other port (“opt-out”). Wait times were defined as how long rats waited before opting out. Trial initiation times were defined as the time from opting-out or consuming reward to initiating a new trial. Reward delays were drawn from an exponential distribution, and on 15-25 percent of trials, rewards were withheld to force rats to opt-out, providing a continuous behavioral readout of subjective value (Fig. 1b)^16–18^. We used a high-throughput facility to train 291 rats using computerized, semi-automated procedures. The facility generated statistically powerful datasets (median = 33,493 behavioral trials, 71 sessions).

**Figure 1:**
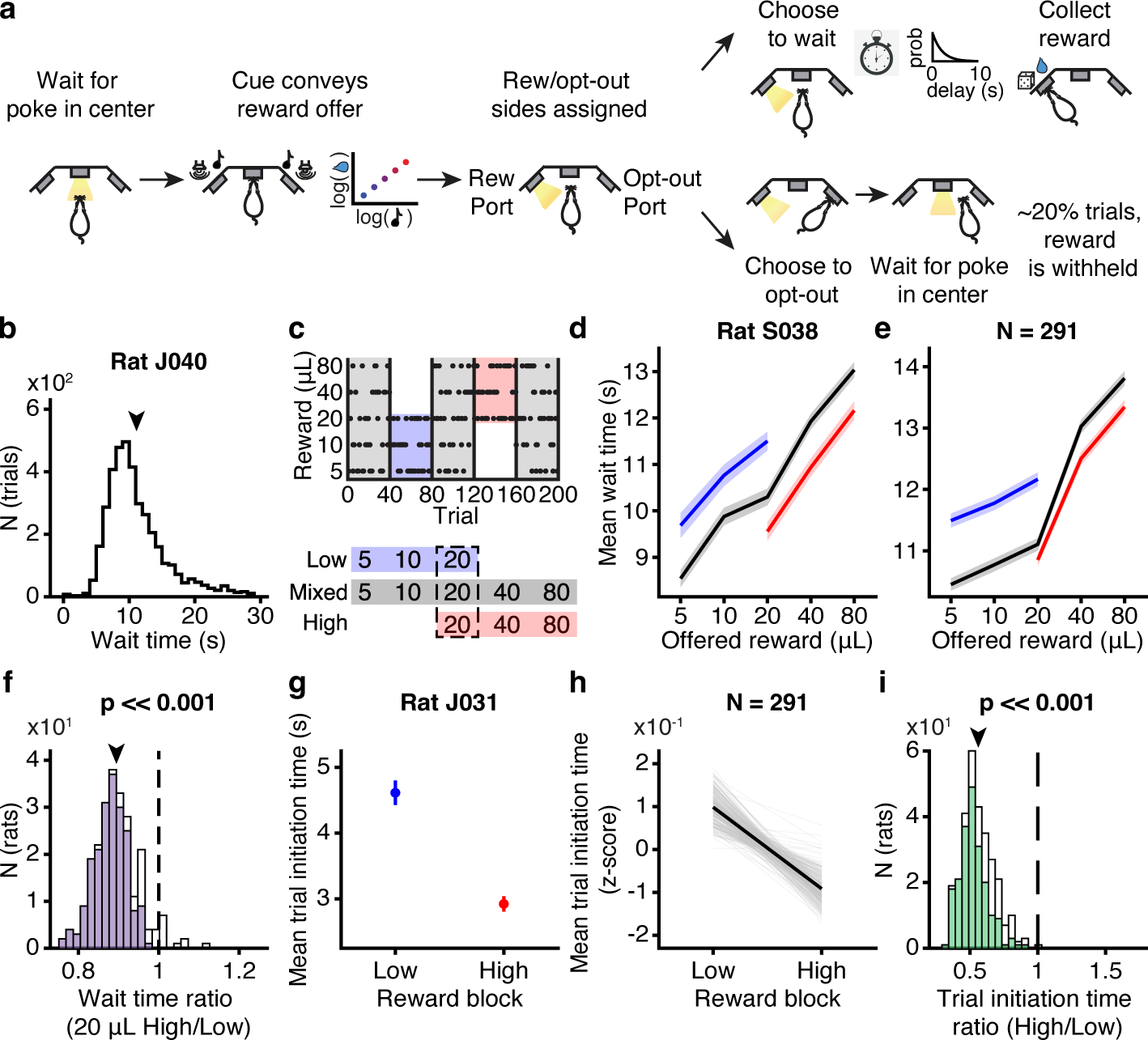
Wait time and trial initiation time were modulated by the value of the environment. **a.** Schematic of behavioral paradigm. **b.** Distribution of wait times for one rat. **c.** Block structure of task. **d-e.** Average wait time on catch trials by reward in each block for (d) one rat and (e) averaged across rats. **f.** Wait time ratio (average wait time for 20 *µ*L in high block/low block) across all rats. Filled boxes indicated rats with *p <* 0.05, Wilcoxon rank-sum test. Population average, *p <<* 0.001, Wilcoxon signed-rank test, *N* = 291. **g-h.** Average trial initiation times in high and low blocks for (g) one rat and (h) all rats. **i.** Trial initiation time ratio (average initiation time in high block/low block) across all rats. Filled boxes indicated rats with *p <* 0.05, Wilcoxon rank-sum test. Population average, *p <<* 0.001, Wilcoxon signed-rank test, *N* = 291. tual effects (i.e., effects of hidden states). The hidden states differed in their average reward and therefore in their opportunity costs, or what the rat might miss out on by continuing to wait. According to foraging theories, the opportunity cost is the long-run average reward, or the value of the environment^19^. In accordance with these theories^19, 20^, rats adjusted how long they were willing to wait for rewards in each block, and on average waited *∼*10 percent less time for 20µL in high blocks, when the opportunity cost was high, compared to in low blocks (*p <<* 0.001, Wilcoxon signed-rank test, *N* = 291; Fig. 1d-f). These are strong contextual effects compared to previous studies^17, 21^.

The task contained latent structure: rats experienced blocks of 40 completed trials (hidden states) in which they were presented with low (5, 10, or 20µL) or high (20, 40, or 80µL) rewards^17^. These were interleaved with “mixed” blocks which offered all rewards (Fig. 1c). 20µL was present in all blocks, so comparing behavior on trials offering this reward revealed contex-

Animals make more vigorous actions when those actions are expected to yield larger or more valuable rewards^2, 22–25^. Therefore, we analyzed how quickly rats initiated trials, as this might also reflect the perceived value of the environment. Indeed, trial initiation times were modulated by blocks in a similar pattern as the wait times, with rats initiating trials more quickly in high compared to low blocks (*p <<* 0.001, Wilcoxon signed-rank test, *N* = 291; Fig. 1g-i; Extended Data Fig. 1). Previous work suggests that this pattern optimally balances the energetic costs of vigor against the benefits of harvesting reward in environments with different reward rates^2, 25, 26^. Therefore, both the trial initiation times, which reflect motivation, and the wait times, which reflect deliberating between waiting and opting-out, were modulated by the value of the environment.

Notably, while we used all behavioral trials for analyses of initiation times in this study, sensitivity to the reward blocks was largely driven by initiation times following unrewarded trials (Extended Data Fig. 2, Methods), which accounted for more variance in initiation times. This is consistent with previous studies showing that response outcomes can gate behavioral strategies^27, 28^. There were no major differences in wait times following rewarded or unrewarded trials (Extended Data Fig. 3). To make comparisons between trial initiation and wait times with as much statistical power as possible and the fewest assumptions, we used all behavioral trials for subsequent analyses in this study.

**Figure 2:**
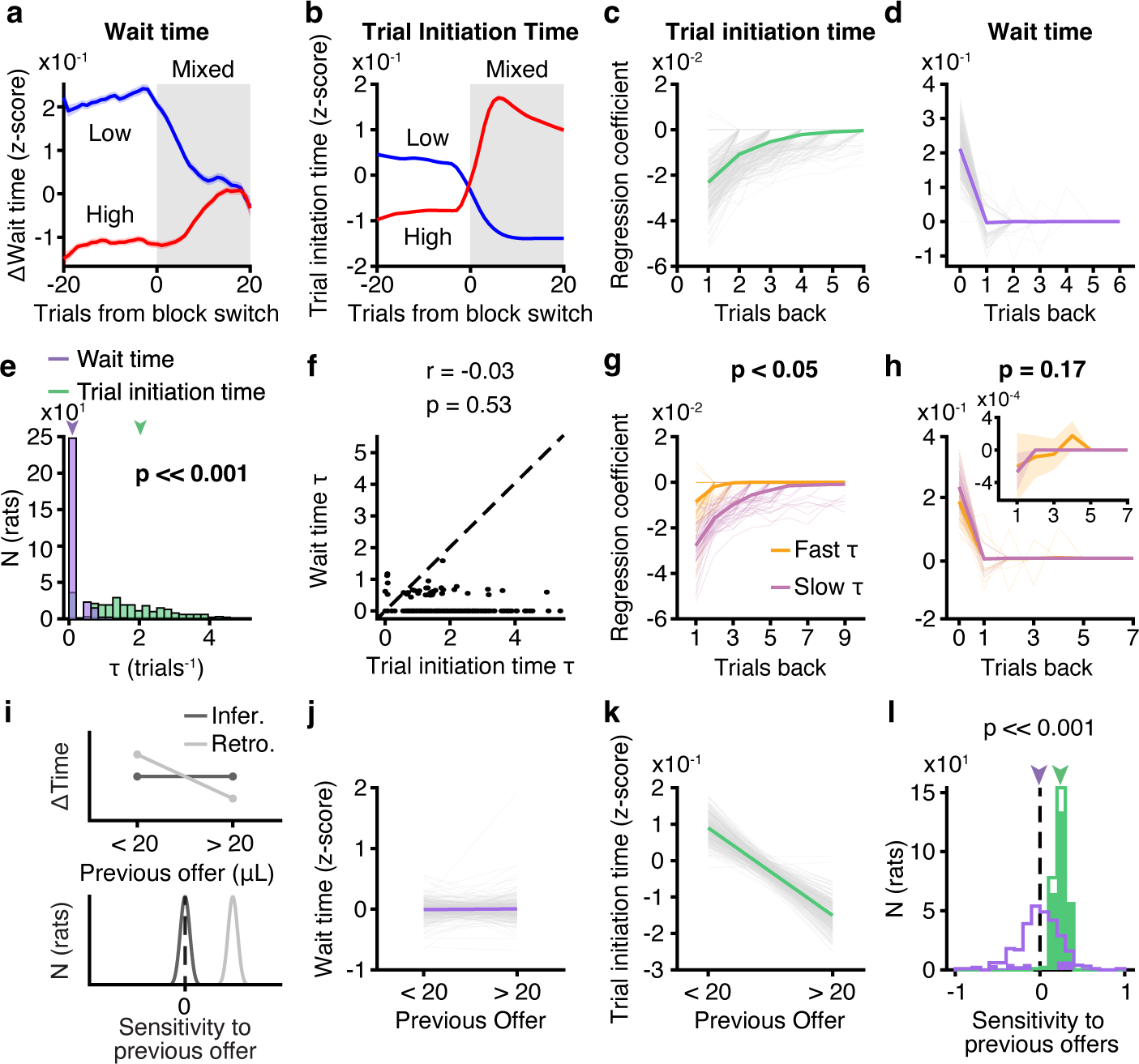
Wait and trial initiation times use distinct estimates of the value of the environ-ment. **a-b.** Mean change in wait times (a) and trial initiation times (b) from low or high blocks to mixed blocks, *N* = 291. Data are mean *±* S.E.M. Data were smoothed with a moving win-dow of 10 trials. **c-d.** Regression coefficients for (c) trial initiation time and (d) wait time. **e-f.** Time constants, *τ*, of exponential decay parameters fit to previous trial coefficients for wait time (purple) and trial initiation time (green) were (e) significantly different, *p <<* 0.001, Wilcoxon sign-rank test, *N* = 291, and (f) uncorrelated, *r* = −0.03, *p* = 0.53, Pearson linear correlation, *N* = 291. **g-h.** Fast or slow initiation time *τ* (*<*20^th^ or *>*80^th^ percentile) meaningfully divided rats based on their initiation time regression coefficients (g; *p <<* 0.01, one-tailed permutation test, *N* = 116), but not wait time coefficients (h; *p* = 0.1, one-tailed permutation test, *N* = 116). Inset shows previous trial coefficients for wait times with adjusted y-axis limits. **i.** Predictions for sensitivity to previous offers (behavior conditioned on previous offer *<*20*µ*L -*>*20*µ*L) for fixed (light) versus sequentially-updated (dark) estimates of environmental value, consistent with in-ferential and retrospective strategies, respectively. **j.** Wait time on 20 *µ*L catch trials in mixed blocks conditioned on previous reward offer. (*p <* 0.05 for 38/291 rats, Wilcoxon rank-sum test). **k.** Trial initiation time in mixed blocks conditioned on previous reward offer. (*p <* 0.05 for 256/291 rats, Wilcoxon rank-sum test). **l.** Sensitivity to previous offers for wait time (pur-ple) and trial initiation time (green). *p <<* 0.001, Wilcoxon sign-rank test, *N* = 291. Colored bars are individual rats with *p <* 0.05, Wilcoxon rank-sum test.

**Figure 3:**
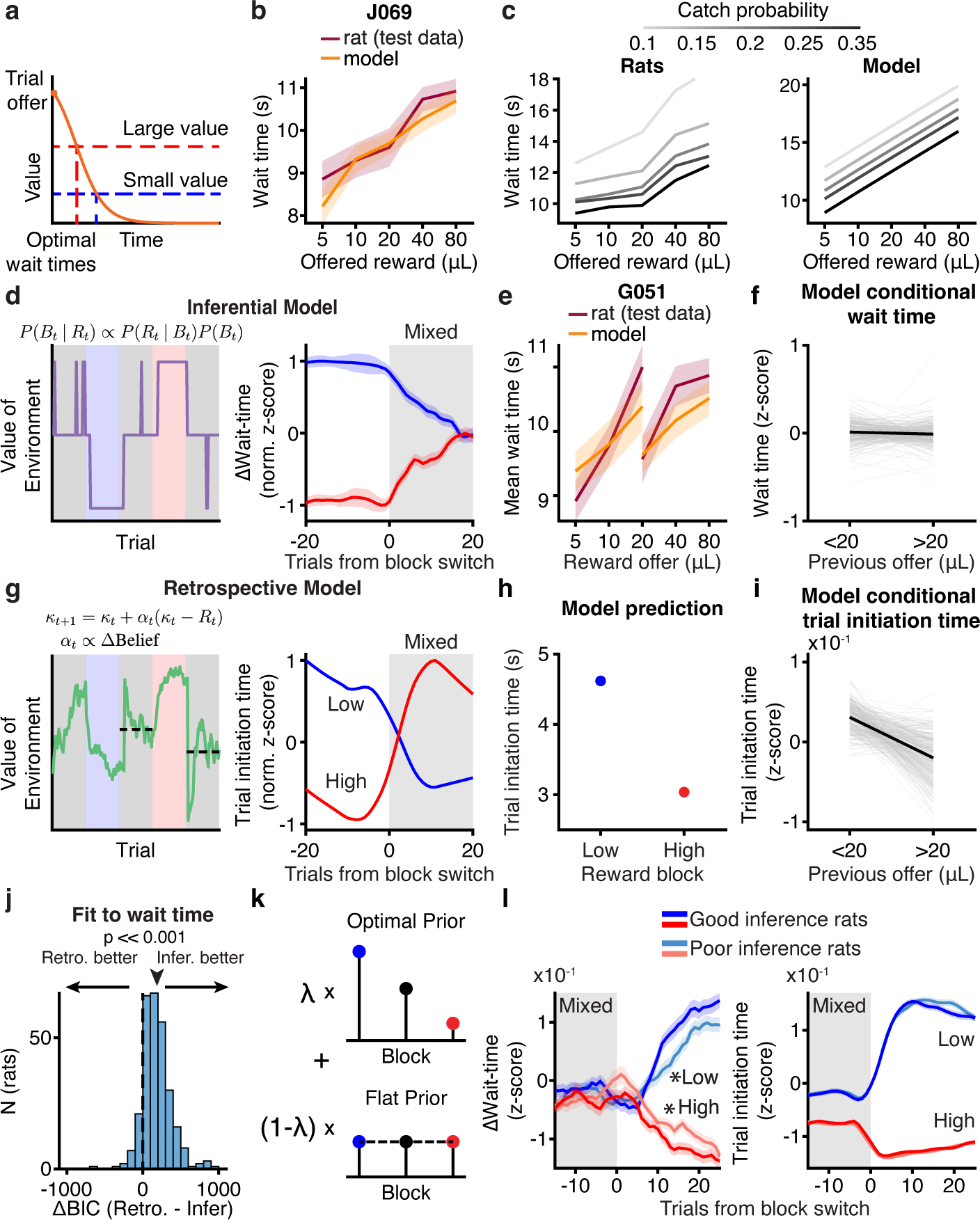
Computational modeling reveals distinct value computations for wait time and trial initiation. **a.** Model schematic. **b.** Example wait time model performance for mixed blocks only in held-out test data. **c.** Rat population (left; sample sizes in methods) and model (right) wait time data in mixed blocks as a function of catch probability. **d.** Example opportunity cost and wait time dynamics from inferential model. **e.** Inferential model fit to rats can capture wait time behavior in held-out test data. **f.** Inferential model captures conditional wait time trend across rats (*N* = 291). **g.** Example opportunity cost and wait time dynamics from retrospective model. **h.** Retrospective model can qualitatively capture trial initiation time behavior. **i.** Retro-spective model captures conditional trial initiation time trend across rats (*N* = 291). **j.** Model comparison using Δ BIC prefers inferential model compared to retrospective model when fit to wait time data (*p <<* 0.001, Wilcoxon Signed-rank test, *N* = 291). **k.** Schematic for sub-optimal inference model **l.** Transitions from mixed to low (blue) or high (red) blocks for wait time (left) or trial initiation time (right) separated by quality of inference (*λ <* 20th or *>* 80th percentile). *p *<* 0.05, one-tailed non-parametric shuffle test comparing logistic fit parameters, N = 116. Data are mean *±* S.E.M.

### Trial initiation and wait times exhibited distinct temporal dynamics

Surprisingly, wait and trial initiation times exhibited dramatically different dynamics at block transitions. In mixed blocks, the wait times following high and low blocks converged to a common value, regardless of the previous block type, suggesting the use of a fixed esti-mate of environmental value in mixed blocks (Fig. 2a). Trial initiation times, however, showed longer timescale effects such that initiation times in mixed blocks strongly depended on the previous block identity (Fig. 2b; Extended Data Fig. 4). These longer timescale dynamics, which are reminiscent of incentive contrast effects^29^, were also evident in the transitions from mixed blocks into high/low blocks for trial initiation times, but not wait times (Extended Data Fig. 5), indicating that trial initiation and wait times utilize distinct estimates of the value of the environment.

**Figure 4:**
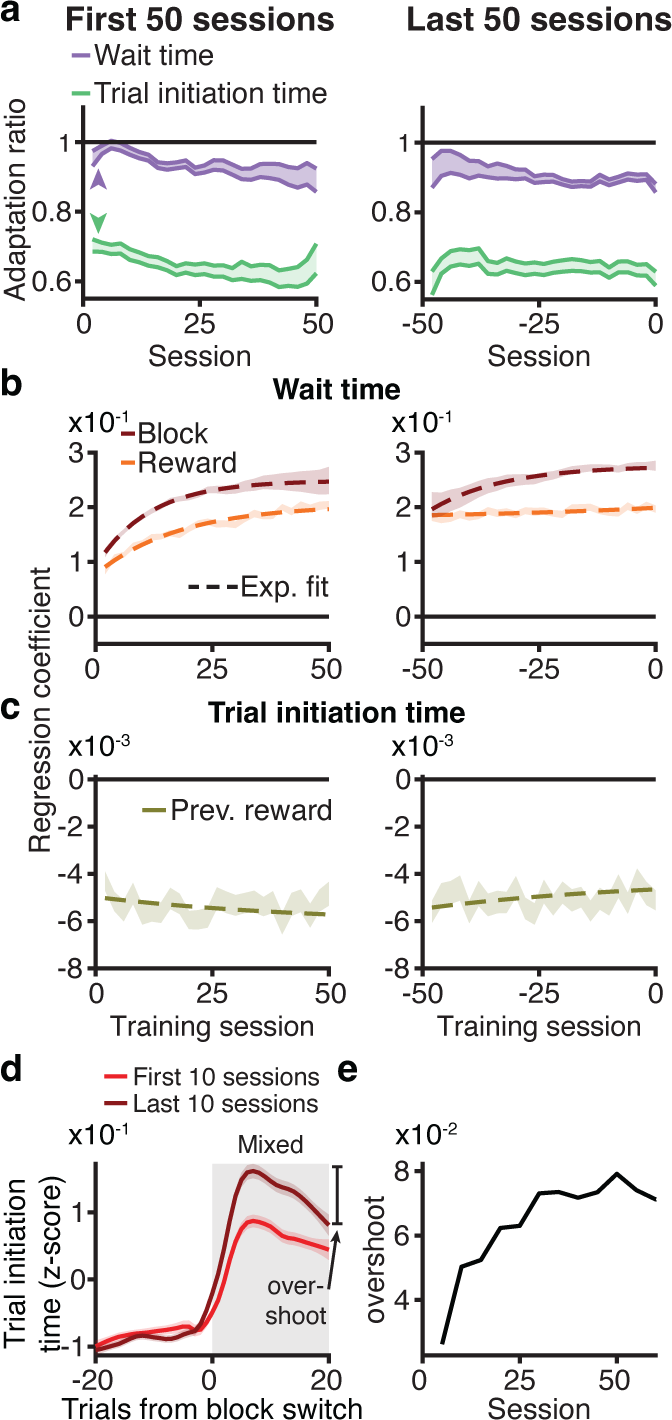
Block sensitivity for wait times requires structure learning. **a.** Wait time adap-tation ratio (average wait time for 20 *µ*L in high/low blocks) evolved over training, while trial initiation time ratio (average in high/low blocks) was below 1 on first session. **b.** Linear regres-sion coefficients for block and reward gradually evolved over training for wait time. **c.** Linear regression coefficient for previous reward was relatively stable across training for trial initiation time. **d.** Overshoot in trial initiation time (difference between maximum z-scored trial initia-tion time and trial initiation time at trial 20 post-transition) was more prominent after structure learning. **e.** Overshoot in trial initiation time dynamics evolved on a similar timescale as block sensitivity for wait times.

**Figure 5:**
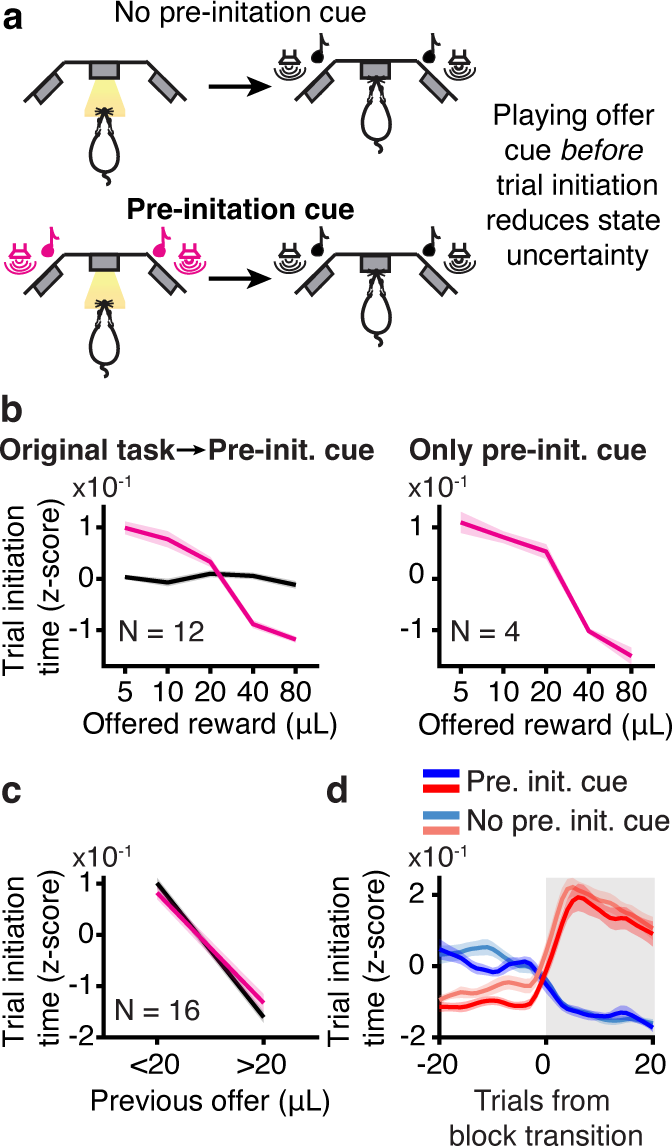
Value computations for motivation do not depend on state uncertainty. **a.** Schematic of pre-initiation cue experiment. **b.** Trial initiation time varied as a function of offered volume for rats that trained on the original task before transitioning to pre-initiation cue task (left) and for rats that trained exclusively on the pre-initiation cue task (right). **c.** Trial initi-ation times were still sensitive to previous reward (behavior on trials offering 20µL conditioned on the previous reward offer) after training on the pre-initiation cue task. (*p <* 0.05 for 13/16 rats, Wilcoxon Rank-sum test, *N* = 16). **d.** Trial initiation times in mixed blocks depended on previous block type in pre-initiation cue task.

To better characterize their temporal dynamics, we regressed the trial initiation and wait times against rewards offered on previous trials. We included current rewards as regressors in the wait time model, and restricted this analysis to mixed blocks only. Examination of the regression coefficients revealed qualitatively different dynamics, in which the wait times were explained by the reward offered on the current trial, but the trial initiation times reflected an exponentially weighted effect of previous rewards, consistent with a model-free temporal dif-ference learning rule (Fig. 2c,d). We fit exponential curves to the previous trial coefficients for each rat, and found that the distributions of exponential decay time constant parameters (*τ*) were significantly different for the trial initiation and wait times (*p <<* 0.01, Wilcoxon sign-rank test, *N* = 291; Fig. 2e). Moreover, *τ* parameters were not correlated across models (*r* = 0.08, *p* = 0.18, Pearson linear correlation, *N* = 291, Fig. 2f).

To leverage individual variability across rats, we compared rats with fast and slow temporal integration for trial initiation times (*τ* from exponential fit to regression coefficients *<* 20th or *>* 80th percentiles). There were differences in temporal integration for trial initiation times, but not wait times, for these groups (Fig. 2g-h, trial initiation time *p <<* 0.001, wait time *p* = 0.5, permutation test, *N* = 116). Collectively, these data suggest that within a block, wait times use a fixed estimate of the value of the environment, whereas trial initiation times are sensitive to previous rewards (Fig. 2c,d). Indeed, for almost all rats (87%), wait times for 20µL offers in mixed blocks were not significantly different if they were preceded by rewards that were smaller or larger than 20µL (*p >* 0.05, Wilcoxon rank-sum test, *N* = 253/291). However, for 89% of rats, trial initiation times were significantly modulated by previous rewards, suggesting fixed and incrementally updated estimates of the value of the environment, respectively (*p <* 0.05, Wilcoxon rank-sum test, *N* = 256/291, Fig. 2i-l).

One factor that could in principle influence initiation times is satiety. However, satiety effects were small, and modestly apparent as a gradual increase in trial initiation times over the session. To control for these effects, we regressed out initiation times against trial number in order to detrend the slow changes over the course of a session. However, there was no qualitative change in any of our results, including the dynamics at block transitions, if we did not detrend (Extended Data Fig. 6). Because the trials are self-initiated, we suspect that when rats are sated they choose not to initiate trials, thereby minimizing the effects of satiety on behavior, at least compared to other factors that contribute more to the variance in initiation times (e.g., reward history, Fig. 2c).

### Computational modeling reveals distinct value computations for sequential decisions

Our data suggest that rats’ sequential decisions (when to initiate trials and how long to wait for rewards) reflect different value computations. We developed behavioral models for wait and trial initiation times, inspired by foraging theories^19^. The wait time model implemented a trial value function that scaled with the offered reward and decayed to reflect reward probability over time^16^. The model’s predicted wait time was when the value function fell below the value of the environment (opportunity cost) on each trial (Fig. 3a). This model captured key features of the rats’ behavior, including the monotonic relationship between wait time and reward offer in mixed blocks (Fig. 3b) as well as a graded dependence of wait times on the catch probability (Fig. 3c). Different versions of the model estimated the value of the environment using different computations.

Analysis of rats’ trial initiation times suggests that they estimate the value of the environment as a running average of rewards (Fig. 2c)^2, 17, 30^. We refer to this computation as retrospective, as it reflects past experience^31^. Alternatively, rats’ wait times reflected the use of discrete estimates of block value (Fig. 2a,d,j). Therefore, rats might infer the current block^31–35^, and use fixed estimates of block value based on that inference. We refer to this computation as inferential, since it requires hidden state inference.

The inferential model selected the most likely block using Bayes’ Rule with a prior that incorporated reward history and knowledge of the block transition structure. This model recapitulated the rats’ wait times converging to a common value in mixed blocks (Fig. 3d-e). Across rats, the model also captured that wait times for 20µL in mixed blocks were not sensitive to previous rewards (Fig 2j; Fig. 3f). This reflects the model’s use of a fixed estimate of the value of the environment in each block.

In the retrospective case, the value of the environment was estimated as a recency-weighted average of offered rewards according to a temporal-difference learning rule (Fig. 3g). A static learning rate was unable to capture the rats’ behavior (Extended Data Fig. 7). Previous work has shown that animals adjust their learning rates depending on the volatility in the environment, since it is advantageous to learn faster in dynamic environments^36–38^. Therefore, our model scaled the learning rate by the trial-by-trial change in the inferential model’s beliefs about the hidden state (derivative of the posterior, see Methods).

The retrospective model captured several key features of rats’ trial initiation times, which we modeled as inversely proportional to the value of the environment^2^ (Fig. 2g-i). First, with a sufficiently small learning rate (*<*0.1, Fig. 2g), the model integrated reward history on long timescales such that trial initiation times in mixed blocks depended on the previous block identity. Second, the dynamic learning rate captured the rapid behavioral dynamics at block transitions. Finally, integrating over previous trials captured the dependence of trial initiation times on previous rewards in mixed blocks (Fig. 2k, Fig. 3i). We explored versions of the dynamic learning rate that did not reflect inference, including using the unsigned reward prediction error or a running average of reward prediction errors^38^. However, these models could not capture both short and long timescale dynamics at block transitions (Extended Data Fig. 7). This suggests that trial initiation times reflect a retrospective computation that is influenced by subjective belief distributions^36, 37^. In other words, while trial initiation and wait times reflect distinct value computations, those computations interact when states are uncertain via a dynamic learning rate.

We fit the retrospective and inferential models to rats’ wait times. By several model comparison metrics, wait times were better fit by the inferential model that used hidden state inference to select block-specific estimates of the value of the environment (*p <<* 0.001, Wilcoxon signed-rank test, *N* = 291; Fig. 3j, Extended Data Fig. 8), consistent with that model reproducing the wait time dynamics (Fig. 2a,3d). We also used the model to identify trials in mixed blocks where the rats were likely to make mistaken inferences. The rats’ wait times reflected these mistaken inferences, further indicating that their wait times were well-described by the inferential model (*p <<* 0.001, Wilcoxon signed-rank test comparing wait times for 20µL in misinferred high vs. low blocks, *N* = 291; Extended Data Fig. 9).

We also developed a “belief state” model that estimated the value of the environment as the sum of block-specific values weighted by their posterior probabilities. The inferential and belief state models make qualitatively similar predictions about the average wait times. In fact, when the posterior beliefs are stable, which is often the case, the belief state and inferential models are identical, and model comparison did not favor one model over the other (Extended Data Fig. 8).

To leverage individual differences, we turned to the inferential model of wait times. We added a parameter, *λ*, that controlled the extent to which the model used an optimal prior, *λ* = 1, versus an uninformative prior, *λ* = 0 (Fig. 3k; Extended Data Fig. 10). We divided the rats into groups with low or high values of *λ* (*λ <* 20th or *>* 80th percentiles; Extended Data Fig. 11), and compared the parameters of logistic functions fit to the average wait time dynamics for these groups. Rats with optimal and poor inference exhibited significantly different dynamics at transitions from mixed into low or high blocks, indicated by different inverse temperature parameters (mix to low/high, *p <* 0.05, one-tailed permutation test, *N* =180 Fig. 3l). There was no difference in the dynamics of trial initiation times for those same groups of rats (mixed to low: *p* = 0.3, mixed to high: *p* = 0.2, one-tailed permutation test, *N* = 180; Fig. 3l). Therefore, individual differences in trial initiation (Fig. 2g,h) and wait times (Fig. 3l) are dissociable.

### Block sensitivity for wait times requires structure learning

Structure learning is the process of learning the hidden structure of environments, including latent states and transition probabilities between them^39^. If wait and trial initiation times differentially required knowledge of latent task structure, they should exhibit different dynamics over training. In the final stage of training, when rats were introduced to the hidden states, their wait times for 20µL gradually became sensitive to the reward block (Fig. 4a). We observed a gradual increase in the magnitude of reward and block regression coefficients that mirrored the behavioral sensitivity to hidden states (Fig. 4b). In contrast, trial initiation times exhibited block sensitivity on the first session in the final training stage (Fig. 4a). This sensitivity was comparable early and late in training, consistent with animals using previous rewards to a similar extent at these timepoints (Fig. 4c). These data suggest that block sensitivity for wait times, but not trial initiation times, required learned knowledge of hidden task states, and that these decisions reflected computations with distinct learning dynamics.

The modest increase in trial initiation time block sensitivity over training is consistent with the gradual use of a dynamic learning rate that reflected learned knowledge of the blocks. A hallmark of the dynamic learning rate was the “overshoot” after transitions from high to mixed blocks (difference between maximum trial initiation time after transitioning and the trial initiation time 20 trials post-transition; Fig. 2b). The overshoot became more prominent with training (Fig. 4d), on a similar timescale as block sensitivity for wait times (Fig. 4e), suggesting a shared mechanism.

### Reducing state uncertainty did not change trial initiation times

Why would animals use a retrospective computation at trial initiation, but rely on an inferential computation as rats deliberated just 1-2 seconds later? In non-human primates, the decision to initiate trials can also reflect retrospectively computed values that differ from the values governing the subsequent choice^40, 41^. One possibility is that motivation and approach behavior rely on neural circuits that do not support inference^11^. Another possibility is that actions more distal to rewards are more likely to be retrospective, because there are more steps required to men-tally simulate outcomes for forward-looking strategies like planning^42, 43^. According to either hypothesis, the decision of when to initiate a trial is inherently retrospective.

Theoretical work in reinforcement learning has suggested that the brain should select the strategy that is the fastest and most accurate when taking into account uncertainty^8, 9^. Therefore, perhaps trial initiation times are retrospective because the rats’ subjective beliefs about the inferred state have more uncertainty before they hear the reward offer. Model simulations of a Bayes’ optimal observer did show that the reward offer reduced the uncertainty of subjective beliefs about the hidden state (comparing variance of prior to variance of posterior, *p <<* 0.001, Wilcoxon sign-rank test).

To test this hypothesis, we modified the task so that some rats heard the reward cue before they initiated the trial, when the center light turned on; they heard the tone again at trial initiation, as in the standard task (Fig. 5a). Their trial initiation times became sensitive to the offered reward (Fig. 5b). However, trial initiation times for 20µL in mixed blocks were still modulated by the previous reward, consistent with the use of incrementally updated estimates of the value of the environment within a block (*p <* 0.05 for 13/16 rats; Fig. 5c). Moreover, how quickly they initiated trials in mixed blocks continued to depend on the previous block identity (Fig. 5d). These data indicate that there may be something inherently retrospective about the motivational decision to initiate a trial.

## Discussion

We used high-throughput training to collect statistically powerful datasets and leverage individual variability across hundreds of animals. Consistent with previous work, rats adjusted their behavior as we varied the richness of the environment in a way consistent with forag-ing theories^19, 30, 44–46^, and behavioral economic theories of reference dependence^47, 48^. Notably, we found that animals used multiple, parallel computations to estimate the richness of the en-vironment, and rapidly switched between these computations on single trials, indicating that value computations vary on fine timescales (seconds). Our data are consistent with evidence for multiple decision-making systems that rely on distinct neural circuits^10, 12, 49, 50^. While ani-mals’ decisions of how long to wait for rewards relied on hidden state inference, the decision of when to initiate the trial was governed by a retrospective computation that calculated the value of the environment as the running average of rewards. Reducing state uncertainty before the trial did not change the value computations governing trial initiation times, suggesting that this decision may be inherently retrospective, although influenced by subjective belief distributions via a dynamic learning rate.

Recent work in psychology, machine learning, and neuroscience has characterized how par-allel value computations might be combined^4–8, 40, 51–54^. For instance, in multi-step decision tasks, interaction effects in regression models are thought to reflect the use of combined ret-rospective and inferential value estimates^14, 15^, and hybrid strategies for computing values have been approximated as a weighted average of retrospective and inference-based values^40^. Our findings add to this body of work. Instead of simply combining or averaging values that were computed in different ways, rats seemed to coordinate their dynamics: changes in subjective beliefs about inferred states acted as a gain on retrospective value learning rates. Moreover, we tested the prevailing hypothesis about arbitration between these parallel value computations, namely, that agents should use the value estimate with the lowest uncertainty^8, 9^. We reduced state uncertainty by playing the reward cue before rats initiated trials. However, trial initiation times still reflected retrospective value computations (Fig. 5c-d). We hypothesize that different neural circuits mediate these rapid sequential decisions (starting the trial versus deciding how long to wait), and that these circuits support or favor distinct value computations due to their connectivity and other neurobiological constraints.

Alternatively, previous work has suggested that actions more distal to rewards are more likely to be retrospective, because there are more steps required to mentally simulate outcomes for forward-looking strategies like planning^42, 43^. Therefore, one potential reason that trial initi-ation times were retrospective is because they were more distal to rewards. However, in multi-step decision-making tasks (i.e., the two-step task), the first action, which is diagnostic of how value is computed, generally reflects computations that use a model of the world to flexibly esti-mate values^14, 55, 56^. Compared to the two-step task, the first action in our task is a similar number of states away from the terminal reward state, but the temporal delays are longer. Therefore, it is possible that temporal proximity to reward may determine how values are computed.

It may be counterintuitive that the retrospective computation produced faster dynamics at block transitions than hidden state inference (Fig. 2a,b). Two features of the models explain this observation. First, the inferential model selects the block with the maximum posterior probability. This argmax operation nonlinearly thresholds whether changes in the posterior produce changes in the inferred state. In contrast, the retrospective model’s estimate of the value of the environment is directly influenced by graded, “subthreshold” changes in the posterior via the dynamic learning rate. Subthreshold changes in the posterior necessarily precede changes that cross threshold for inferring a state change. Second, the inferential model’s prior is recursive: the posterior on one trial becomes the prior on the next trial. This means that the prior accumulates information over trials to infer state changes, instead of making them instantaneously. Indeed, individual differences in the informativeness of rats’ priors predicted the dynamics of their inferred state changes (Fig. 3l).

The contextual effects we observed likely reflect efficient coding of value^17, 57–59^. According to the efficient coding hypothesis, to represent stimuli efficiently, neurons should be tuned to stimulus distributions that animals are most likely to encounter in the world^60^. Recent studies have shown that biases in value-based decision-making, including the contextual effects observed here, reflect efficient value coding^17, 57, 58^. Previous studies examined how neurons “adapted” to reward or stimulus distributions over blocks of trials or sessions, implying gradual, experience-dependent adjustments in behavioral sensitivity and neural tuning^17, 61, 62^. Our findings suggest that if animals have learned the reward or stimulus distributions associated with a particular state, they can condition their subjective value representations on that inferred state, perhaps via discrete, state-dependent adjustments in neural sensitivity^63^. A major future question is how multi-regional neural circuits represent belief distributions for hidden state inference, and condition rapid adjustments in efficient neural representations of value on inferred states.

## Methods

### Subjects

A total of 291 Long-evans rats (184 male, 107 female) between the ages of 6 and 24 months were used for this study (*Rattus norvegicus*). The Long-evans cohort also included ADORA2A-Cre (*N* =10), ChAT-Cre (*N* =2), DRD1-Cre (N=3), and TH-Cre (*N* =12). Animal use procedures were approved by the New York University Animal Welfare Committee (UAWC #2021-1120) and carried out in accordance with National Institutes of Health standards.

Rats were pair housed when possible, but were occasionally single housed (e.g. if fighting occurred between cagemates). Animals were water restricted to motivate them to perform be-havioral trials. From Monday to Friday, they obtained water during behavioral training sessions, which were typically 90 minutes per day, and a subsequent ad libitum period of 20 minutes. Following training on Friday until mid-day Sunday, they received ad libitum water. Rats were weighed daily.

### Behavioral training

Rats were trained in a high-throughput behavioral facility in the Constantinople lab using a computerized training protocol. They were trained in custom operant training boxes with three nose ports. Each nose port was 3-D printed, and the face was protected with an epoxied stainless steel washer (McMaster-Carr #92141A056). All ports contained a visible light emit-ting diode (LED; Digikey #160-1850-ND), and an infrared LED (Digikey #365-1042-ND) and infrared photodetector (Digikey #365-1615-ND) that enabled detection of when a rat broke the infrared beam with its nose. Additionally, the side ports contained stainless steel lick tubes (McMaster-Carr #8988K35, cut to 1.5mm) that delivered water via solenoid valves (Lee Com-pany #LHDA1231115H). There was a speaker mounted above each side port that enabled de-livery of stereo sounds (Bohlender Graebener). The behavioral task was instantiated as a finite state machine on an Arduino-based behavioral system with a Matlab interface (Bpod State Machine r2, Sanworks), and sounds were delivered using a low-latency analog output module (Analog Output Module 4ch, Sanworks) and stereo amplifier (Lepai LP-2020TI).

Research technicians loaded rats in and out of the training rigs in each session, but the training itself was computer automated. All rig computers automatically pulled version-controlled software from a git repository and wrote behavioral data to a MySQL (MariaDB) database hosted on a synology server. Rig computers automatically loaded each rat’s training settings file from the previous session, and following training, wrote a new settings file to the server for the subsequent day of training. Rig computers automatically loaded files for specific rats based on a schedule on the MySQL database. Human intervention was possible but generally unnecessary.

### Sound Calibration

We calibrated sounds using a hand-held Precision Sound Level Meter with a 1/2” microphone (Bruel & Kjaer, Type 2250). The microphone was calibrated with a sound level calibrator (Bruel & Kjaer, Type 4230). Tones of different frequencies (1, 2, 4, 8, 16kHz) were presented for 10 seconds each; these tones were selected because they are in the trough of the behavioral audiogram for rats^64^. They are also on a logarithmic scale and thus should be equally discriminable to the animals. We adjusted the auditory gain in software for each frequency stimulus to match the sound pressure level to 70dB in the rig, measured when the microphone was proximal to the center poke.

### Task Logic

LED illumination from the center port indicated that the animal could initiate a trial by poking its nose in that port - upon trial initiation the center LED turned off. While in the center port, rats needed to maintain center fixation for a duration drawn uniformly from [0.8, 1.2] seconds. During the fixation period, a tone played from both speakers, the frequency of which indicated the volume of the offered water reward for that trial [1, 2, 4, 8, 16kHz, indicating 5, 10, 20, 40, 80*µ*L rewards]. Following the fixation period, one of the two side LEDs was illuminated, indicating that the reward might be delivered at that port; the side was randomly chosen on each trial. This event (side LED ON) also initiated a variable and unpredictable delay period, which was randomly drawn from an exponential distribution with mean = 2.5 seconds. The reward port LED remained illuminated for the duration of the delay period, and rats were not required to maintain fixation during this period, although they tended to fixate in the reward port. When reward was available, the reward port LED turned off, and rats could collect the offered reward by nose poking in that port. The rat could also choose to terminate the trial (opt-out) at any time by nose poking in the opposite, un-illuminated side port, after which a new trial would immediately begin. On a proportion of trials (15-25%), the delay period would only end if the rat opted out (catch trials). If rats did not opt-out within 100s on catch trials, the trial would terminate.

The trials were self-paced: after receiving their reward or opting out, rats were free to initiate another trial immediately. However, if rats terminated center fixation prematurely, they were penalized with a white noise sound and a time out penalty (typically 2 seconds, although adjusted to individual animals). Following premature fixation breaks, the rats received the same offered reward, in order to disincentivize premature terminations for small volume offers.

We introduced semi-observable, hidden-states in the task by including uncued blocks of trials with varying reward statistics^17^: high and low blocks, which offered the highest three or lowest three rewards, respectively, and were interspersed with mixed blocks, which offered all volumes. There was a hierarchical structure to the blocks, such that high and low blocks alternated after mixed blocks (e.g., mixed-high-mixed-low, or mixed-low-mixed-high). The first block of each session was a mixed block. Blocks transitioned after 40 successfully completed trials. Because rats prematurely broke fixation on a subset of trials, in practice, block durations were variable.

### Criteria for including behavioral data

In this task, the rats were required to reveal their subjective value of different reward offers. To determine when rats were sufficiently trained to understand the mapping between the auditory cues and water rewards, we evaluated their wait time on catch trials as a function of offered rewards. For each training session, we first removed wait times that were greater than two standard deviations above the mean wait time on catch trials in order to remove potential lapses in attention during the delay period (this threshold was only applied to single sessions to determine whether to include them). Next, we regressed wait time against offered reward and included sessions with significantly positive slopes that immediately preceded at least one other session with a positive slope as well. Once performance surpassed this threshold, it was typically stable across months. Occasional days with poor performance, which often reflected hardware malfunctions or other anomalies, were excluded from analysis. We emphasize that the criteria for including sessions in analysis did not evaluate rats’ sensitivity to the reward blocks. Additionally, we excluded trial initiation times above the 99th percentile of the rat’s cumulative trial initiation time distribution pooled over sessions.

### Shaping

The shaping procedure was divided into 8 stages. For stage 1, rats learned to maintain a nose poke in the center port, after which a 20 *µ*L reward volume was delivered from a random illuminated side port with no delay. Initially, rats needed to maintain a 5 ms center poke. The center poke time was incremented by 1 ms following each successful trial until the center poke time reached 1 s, after which the rat moved to stage 2.

Stages 2-5 progressively introduced the full set of reward volumes and corresponding auditory cues. Rats continued to receive deterministic rewards with no delay after maintaining a 1 second center poke. Each stage added one additional reward that could be selected on each trial-stage 2 added 40 *µ*L, stage 3 added 5 *µ*L, stage 4 added 80 *µ*L, and stage 5 added 10 *µ*L. Each stage progressed after 400 successfully completed trials. All subsequent stages used all 5 reward volumes.

Stage 6 introduced variable center poke times, uniformly drawn from [0.8-1.2] s. Additionally, stage 6 introduced deterministic reward delays. Initially, rewards were delivered after a 0.1 s delay, which was incremented by 2 ms after each successful trial. After the rat reached delays between 0.5 and 0.8 s, the reward delay was incremented by 5 ms following successful trials. Delays between 0.8 and 1 s were incremented by 10 ms, and delays between 1 and 1.5 s were incremented by 25 ms. Rats progressed to stage 7 after reaching a reward delay of 1.5 s.

In stage 7, rats experienced variable delays, drawn from an exponential distribution with mean of 2.5 seconds. Additionally, we introduced catch trials (see above), with a catch probability of 15%. Stage 7 terminated after 250 successfully completed trials.

Finally, stage 8 introduced the block structure (see above). We additionally increased the catch probably for the first 1000 trials to 35%, to encourage the rats to learn that they could opt-out of the trial. After 1000 completed trials, the catch probability was reduced to 15-20%. All data in this paper was from training stage 8.

### Training for male and female rats

We collected data from both male and female rats (160 male, 114 female). Male and female rats were trained in identical behavioral rigs with the same shaping procedure described above. Early cohorts of female rats experienced the same reward set as the males. However, female rats are smaller, and they consumed less water and performed substantially fewer trials than the males. Therefore, to obtain sufficient behavioral trials from them, reward offers for female rats were slightly reduced while maintaining the logarithmic spacing: [4, 8, 16, 32, 64 *µ*L]. For behavioral analysis, reward volumes were treated as equivalent to the corresponding volume for the male rats (e.g., 16 *µ*L trials for female rats were treated the same as 20 *µ*L trials for male rats). The auditory tones were identical to those used for male rats. We did not observe any significant differences between the male and female rats, in terms of contextual effects (Extended Data Fig. 12), or behavioral dynamics at block transitions (data not shown).

| Stage | Center poke time | 5μL | 10μL | 20μL | 40μL | 80μL | Reward delay | Reward probability | Blocks |
| --- | --- | --- | --- | --- | --- | --- | --- | --- | --- |
| 1 | Increment to 1s |  |  | **X** |  |  | 0 | 1 |  |
| 2 | 1s |  |  | **X** | **X** |  | 0 | 1 |  |
| 3 | 1s | **X** |  | **X** | **X** |  | 0 | 1 |  |
| 4 | 1s | **X** |  | **X** | **X** | **X** | 0 | 1 |  |
| 5 | 1s | **X** | **X** | **X** | **X** | **X** | 0 | 1 |  |
| 6 | Variable (0.8-1.2s) | **X** | **X** | **X** | **X** | **X** | Increment to 1.5s | 1 |  |
| 7 | Variable (0.8-1.2s) | **X** | **X** | **X** | **X** | **X** | Variable (from exponential) | 0.85 |  |
| 8 | Variable (0.8-1.2s) | **X** | **X** | **X** | **X** | **X** | Variable (from exponential) | 0.65-0.85 | **X** |

We tracked most female rats’ stages in the estrus cycle using vaginal cytology, with vaginal swabs collected immediately after each session using a cotton-tipped applicator first dipped in saline. Samples were smeared onto a clean glass slide and visually classified under a light microscope. For the current study, data from female rats was averaged across all stages of the estrus cycle.

### Behavioral models

We developed separate behavioral models to describe rat’s wait time and trial initiation time data. Both wait time and trial initiation time should depend on the value of the environment. For the wait time data, we adapted a model from^16^ which described the wait time, WT, in terms of the value of the environment (i.e., the opportunity cost), the delay distribution, and the catch probability (i.e., the probability of the trial being unrewarded). Given an exponential delay distribution, we defined the predicted wait time as

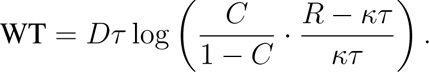

where *τ* is the time constant of the exponential delay distribution, *C* is the probability of reward (1-catch probability), *R* is the reward on that trial, *κ* is the opportunity cost, and *D* is a scaling parameter. In the context of optimal foraging theory and the marginal value theorem, which provided the theoretical foundation for this model, each trial is a depleting “patch” whose value decreases as the rat waits^19^. Within a patch, the decision to leave depends on the overall value of the environment, *κ*, which is stable within trials but can vary across trials and hidden reward states, i.e., blocks.

The above equation was shown to be normative for a Markov Decision Process in which the value of the environment was constant for the foreseeable future^16^. However, given that the value of the environment changed over blocks in our task, it is possible that this equation is not normative for our case. However, this formulation qualitatively captured features of the data, including the graded dependence of wait times on the catch probability, and sensitivity to reward volumes and blocks. Therefore, we found it to be a useful, if not necessarily normative, process model of behavior.

For the trial initiation time, we adapted a model^2^ which describes the optimal trial initiation time, TI, given the value of the environment, *κ*, as

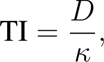

where *D* is a scale parameter.

We initially evaluated two different ways of calculating the value of the environment for these models, which are shared between the wait time and trial initiation time models: a retrospective and inferential model (see below). We assumed independent log-normal noise for each trial, with a constant variance of 8 seconds for the wait time model and 4 seconds for the trial initiation time model. The log-normal noise model outperformed alternative noise models, such as gamma and ex-Gaussian noise. The noise variance terms were selected from a grid search using data from a subset of animals.

### Inferential model

The inferential model has three discrete value parameters (*κ*_low_*, κ*_mixed_*, κ*_high_), each associ-ated with a block. For each trial, the model chooses the *κ* associated with the most probable block given the rat’s reward history. Specifically, for each trial, Bayes’ Theorem specifies the following:

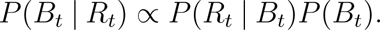

where *B_t_* is the block on trial *t* and *R_t_* is the reward on trial *t*. The likelihood, *P* (*R_t_ | B_t_*), is the probability of the reward for each block, for example,

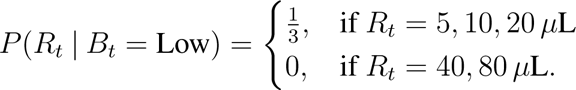

To calculate the prior over blocks, *P* (*B_t_*), we marginalize over the previous block and use the previous estimate of the posterior:

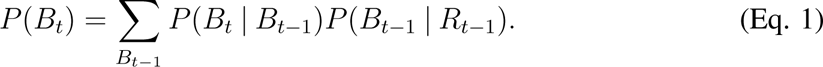

*P* (*B_t_ | B_t−_*_1_), referred to as the “hazard rate,” incorporates knowledge of the task structure, including the block length and block transition probabilities. For example,

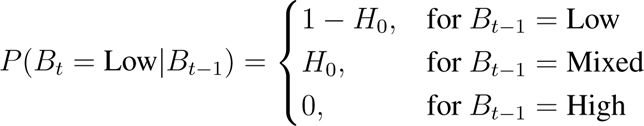

where *H*_0_ = 1*/*40, to reflect the block length. The model assumed a flat block hazard rate for the following reasons. (1) Since animals broke center fixation on a subset of trials, the actual block duration was highly variable. Based on the distributions of experienced block durations, it is unlikely that rats would have learned a perfect step function hazard rate. (2) The blocks spanned several to tens of minutes, making it unlikely that rats would keep a running tally of trials on such long timescales. (3) Gradual changes in wait times at block transitions are not consistent with the use of a veridical step-function hazard rate. (4) We considered an alternative parameterization in which the veridical step function hazard rate was blurred with a Gaussian, but this would have required a number of nontrivial design choices, such as whether the trial counter should be reset after “misinferred” block transitions, regardless of when they occurred in the actual block. (5) Wait times reflected misinferred blocks based on a constant block hazard rate (Extended Data Fig. 9), suggesting that this simplification was a reasonable approximation of the inference process. Including *H*_0_ as an additional free parameter did not improve the performance of the wait time model evaluated on held-out test data in a subset of rats (data not shown), so *H*_0_ was treated as a constant term.

The model selected a fixed value of the environment associated with the most likely block. This formulation is related to an established approximation for solving partially-observable Markov decision processes (POMDPs) known as the Most Likely State algorithm^65^. This algorithm is well-studied, has precedence in the literature as a heuristic approximation for the full posterior distribution over states, and may be biologically plausible as it is computationally tractable compared to more complex solutions to POMDPs.

### Belief state model

Like the inferential model (above), the belief state model has three distinct value parameters and calculates the probability of being in each block using Bayes Rule. However, rather than selecting a single value associated with the most probable block, the model uses the sum of each value, weighted by that probability, that is,

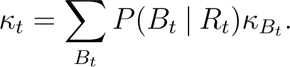

While this model uses the full posterior distribution over states, model comparison found that it was comparable to the simpler Most Likely State algorithm (above; Extended Data Fig. 8). In fact, when the belief distributions were stable (e.g., in adaptation blocks), these models were identical. For that reason, we exclusively used the Most Likely State model (above) for this paper.

There may be alternative normative strategies for this task given different sets of assump-tions. For instance, assuming an infinite time horizon, one might compute the average kappa under the Markov process determining block transitions, starting from the current state. With a sufficiently long time horizon, this average will be dominated by the steady-state distribution of the Markov process, which would predict no contextual modulation of wait times. Given that the rats exhibited strong contextual effects, this strategy is not consistent with their behavior. We therefore did not explore such a model in the current manuscript.

### Inferential model with lambda parameter

To account for potentially sub-optimal inference across rats, we developed a second in-ferential model. This model also uses Bayes rule to calculate the block probabilities, except with a sub-optimal prior, Prior_subopt_. Specifically, we introduce a parameter, *λ*, that generates the sub-optimal prior by weighting between the true, optimal prior (*P* (*B_t_*), Eq. 1), and a flat, uninformative prior (Prior_flat_, uniformly 1*/*3), that is,

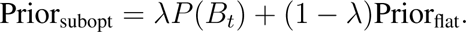

When *λ* = 1, this model reduces to the optimal inferential model, and when *λ* = 0, this model uses a flat prior and the block probabilities are driven by the likelihood.

### Retrospective model

The retrospective model has a single, trial-varying *κ* variable which represents the recency-weighted average of all previous rewards. This average depends on the learning rate parameter *α* with the recursive equation

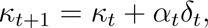

where *κ_t_* is the value of the environment on trial *t*, *r_t_* is the reward on trial *t*, *δ_t_*= *r_t_ − κ_t_* is the reward prediction error (RPE), and *α_t_* is a dynamic learning rate given by *α_t_* = *G · α*_0_. In order to capture the dynamics of the trial initiation times around block transitions, we included a gain term, *G_t_* on the learning rate, which is inversely related to the trial-by-trial change in the mixed block probability from by the inferential model, given by

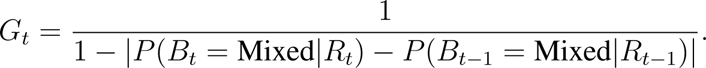

We used trial-by-trial changes in the mixed block probability as a summary statistic of changes in the full posterior distribution. Given the distribution of rewards and the transition structure between blocks, there is always some ambiguity about whether the hidden state is a mixed block, and the posterior block probabilities sum to one. Therefore, changes in the mixed block probability reflect changes in the full posterior on every trial.

The dynamic learning rate we implemented is consistent with previous work showing that humans and animals can adjust their learning rates depending on the volatility and uncertainty in the environment^36–38^. Other models using either (1) a single, static learning rate (*G* = 1), or (2) a dynamic learning rate where the gain term was the unsigned reward prediction error on that trial (*G* = *|δ_t_|*) were unable to capture the observed trial initiation time dynamics at block transitions (Extended Data Fig. 7).

### Fitting and evaluating models

We used MATLAB’s constrained minimization function, fmincon, to minimize the sum of the negative log likelihoods with respect to the model parameters. 100 random seeds were used in the maximum likelihood search for each rat; parameter values with the maximum likelihood of these seeds were deemed the best fit parameters. Before fitting to rat’s data, we confirmed that our fitting procedure was able to recover generative parameters (Extended Data Fig. 13). When evaluating model performance fit to rat data, we performed 5-fold cross-validation and evaluated the predictive power of the model on the held-out test sets. To compare the different models, we used Bayesian Information Criterion (BIC), BIC = log(*n*) *· k* + 2 *·* nLL, where *n* is the number of trials, *k* is the number of parameters, and nLL is the negative log-likelihood of the best-fit model evaluated on all data. We confirmed the model comparison by also comparing Akaike Information Criterion (AIC = 2 *· k* + 2 *·* nLL where *k* is the number of parameters and nLL is the negative log-likelihood of the best-fit model evaluated on all data) and cross-validated negative log-likelihood, which gave similar results to BIC.

We only fit models to the rats’ wait time data. This is because the distribution of trial initiation times was generally heavy-tailed, and seemed to reflect multiple processes on different interacting timescales (e.g., reward sensitivity on short timescales, attention, motivation, and satiety on longer timescales). These processes made it challenging to fit the data with a single process model. Therefore, we used the inferential and retrospective trial initiation time models to generate qualitative predictions that we could compare to the rats’ data.

### Statistical analyses

#### Wait time and trial initiation times: sensitivity to reward blocks

For all analyses, we removed wait times that were one standard deviation above the pooled-session mean. Without thresholding, the contextual effects are qualitatively similar. Outlier wait times tend to occur in low blocks, likely due to attentional or motivational lapses. Therefore, the main difference is that the wait time curves in low blocks are both flatter and longer compared to the thresholded data (Extended Data Fig. 14). When assessing whether a rat’s wait time differed by blocks, we compared each rat’s wait time on catch trials offering 20 *µ*L in high and low blocks using a non-parametric Wilcoxon rank-sum test, given that the wait times are roughly log-normally distributed. We defined each rat’s wait time ratio as the average wait time on 20*µ*L catch trials in high blocks/low blocks. For trial initiation times, we compared all trial initiation times for each block, again using a non-parametric Wilcoxon rank-sum test. We defined each rat’s trial initiation time ratio as the average trial initiation time in high blocks/low blocks.

Trial initiation times were bimodally distributed, with the different modes reflecting whether previous trials were rewarded or not. Unrewarded trials included opt-out trials and trials where rats prematurely terminated center fixation (“violation trials”). Analyzing these trial types separately showed that trial initiation times following unrewarded trials were modulated by blocks in a similar pattern as the wait times, with rats initiating trials more quickly in high compared to low blocks (Extended Data Fig. 2). While we used all behavioral trials for analyses of trial initiation times throughout the manuscript, we note that trial initiation times following rewarded trials exhibited a different pattern (Extended Data Fig. 2), consistent with previous studies showing that response outcomes gate behavioral strategies^27, 28^. Specifically, following rewarded trials, there was a weak positive correlation between reward magnitude and trial initiation time, in contrast to the strong negative correlation we observed following unrewarded trials. We interpret the positive correlation as potentially reflecting micro-satiety effects. However, as these effects were weak, most of the variance in the trial initiation times were driven by those following unrewarded trials.

To assess block effects across the population, we first z-scored each rat’s wait time on all catch trials and trial initiation time on all trials. For wait times, we computed the average z-scored wait time on catch trials offering 20 *µ*L in high and low blocks for each rat, and compared across the population using a paired Wilcoxon sign-rank test. Similarly for trial initiation times, we averaged all z-scored trial initiation times for high and low blocks for each rat, and compared across the population using a paired Wilcoxon sign-rank test.

To assess the effects of catch probability on wait times, we trained cohorts of rats with different catch probabilities. The cohorts varied in size: *N* = [3, 183, 61, 151, 39] for catch probability = [0.1, 0.15, 0.2, 0.25, 0.35], respectively.

#### Block transition dynamics

To examine behavioral dynamics around block transitions, for each rat, we first z-scored wait-times for catch trials of each volume separately in order to control for reward volume effects. We then computed the difference in z-scored wait times for each volume, relative to the average z-scored wait time for that volume, in each time bin (trial relative to block transition), before averaging the differences over all volumes (Δ z-scored wait time). For trial initiation times, we z-scored all trial initiation times. In order to remove satiety effects, for each session individually, we regressed trial initiation time against z-scored trial number and subtracted the fit.

For each transition type, we averaged the Δ z-scored wait times and trial initiation times based on their distance from a block transition, including violation trials (e.g., averaged all wait times four trials before a block transition). Finally, for each block transition type, we smoothed the average curve for each rat using a 10-point moving average, before averaging over rats.

When comparing block transition dynamics in rats with different quality priors, specifically from mixed blocks to high or low, we chose rats in the top or bottom 20^th^ percentile of fit *λ*’s and averaged each group’s block transition dynamics for both wait time and trial initiation time. To compare the normalized dynamics of each group, we fit 4-parameter logistic functions of the following form:

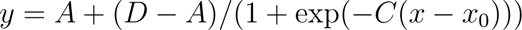

to the behavioral curves and compared the four parameters: *A* (the lower asymptote), *D* (the upper asymptote), *C* (the inverse temperature), and *x*_0_ (*x*-value of the the sigmoid’s midpoint). To determine significance for our observed differences, we performed a non-parametric shuffle test. We generated null distributions on differences in the fit parameters by shuffling the labels of the upper and lower percentile *λ* rats, refitting the logistic to the new shuffled groups’ average dynamic curves, and comparing the fit parameters 500 times. We then used these null distributions to calculate p-values for the observed differences in parameters: the area under this distribution evaluated at the actual difference of parameter values (between high and low *λ* rats) was treated as the p-value.

#### Trial history effects

To assess wait time sensitivity to previous offers, we focused on 20 *µ*L catch trials in mixed blocks only. We z-scored the wait times of these trials separately. Next, we averaged wait times depending on whether the previous offer was greater than or less than 20 *µ*L. For trial initiation times, we used all 20 *µ*L trials in mixed blocks. We averaged z-scored trial initiation times depending on whether the previous offer was greater or less than 20 *µ*L. For both wait time and trial initiation time, we defined the sensitivity to previous offers as the difference between average wait time (trial initiation time) for trials with a previous offer less than 20 *µ*L and trials with a previous offer greater than 20 *µ*L. We compared wait time and trial initiation time sensitivity to previous offers across rats using a paired Wilcoxon signed-rank test.

To capture longer timescale sensitivity across rewards, we regressed previous rewards against wait time and trial initiation time. We focused only on mixed blocks. Additionally, we lin-earized the rewards by taking the binary logarithm of each reward (log_2_(reward)). For wait time, we z-scored wait times for catch trials in mixed blocks. Then, we regressed wait times on these trials against the current offer and previous 9 log_2_(reward) offers, including violation trials, along with a constant offset term. Reward offers from a different block (e.g., a previous high block) were given NaN values. For trial initiation times, we again z-scored for mixed block trials only. Then, we regressed against the previous 9 log_2_(reward) offers, not including the current trial, along with a constant offset. Additionally, we set the reward for violation and catch trials to 0, since rats do not receive a reward on these trials.

For both wait time and trial initiation time, we used Matlab’s builtin regress function to perform the regression. With the coefficients, we found the first non-significant coefficient (coefficient that whose 95% confidence interval contained 0), and set that coefficient and all following coefficients to 0. Finally, we fit a negative exponential decay curve, *y* = *D* exp *−x/τ*, to each rat’s previous trial coefficients (that is, only the previous 9 trial coefficients) for both wait time and trial initiation time and reported the time constant of the exponential decay (tau) for each. If all previous trial coefficients were equal to 0 (as was the case for a vast majority of the wait time coefficients), the time constant was reported as 0. We correlated wait time regression time-constants and trial initiation time regression time-constants using Matlab’s builtin corr function.

#### Learning Dynamics

To assess learning dynamics, we included all sessions after stage 8, not just the sessions that passed criteria for inclusion (above). Because of data limitations examining each session individually (e.g., not every session included both a high and low block), we grouped subsequent sessions into pairs (i.e., we grouped sessions 1 and 2, sessions 3 and 4, etc.). For each session-pair, we calculated the wait time and trial initiation time ratios as above. To assess the emergence of block effects on wait time data, we regressed wait time for each session against both the current reward and a categorical variable representing the current block identity (1 = low block, 2 = mixed block, 3 = high block). To assess the emergence of previous trial effects on trial initiation time, we regressed trial initiation time for each sessions against the previous reward. We smoothed each regression coefficient over sessions using a 5-session moving average. Finally, we set outlier coefficients (3 scaled median absolute deviations away from a 5-point moving median, using Matlab’s builtin *isoutlier* function) to NaN. Finally, we averaged regression coefficients over sessions across rats.

### Pre-initiation cue task

To modulate the subjective uncertainty in the rat’s estimate of state (block) before trial initiation time, we ran a subset of rats on a variation of the task where we cued reward offer before rats initiated a trial (*N* =16). All other aspects of the task remained identical: reward offer cued played again after the rat initiated the trial, rats waited uncued exponentially-distributed delays for rewards, etc. We included both rats that initially trained on the original task before switching to the pre-initiation cue task (*N* = 12), as well as rats who were trained only on the pre-initiation cue task (*N* = 4). To allow the rats who had started on the original task time to adjust to the new task, we only included data after 30 pre-initiation cue sessions. For the rats who were exclusively trained on the pre-initiation cue task, we included all stage 8 sessions. For all rats, we did not exclude sessions using the wait time critera (see above).

To compare effects for rats who had started on the original task, we performed all analyses for data collected on the original task and on the pre-initiation cue task. First, to confirm that the rats learned that the tone before trial initiation indicated the upcoming reward, we averaged z-scored trial initiation times by the offered reward in mixed blocks. We excluded post-violation trials in the original task session, because those trials repeat the same volume as the previous trial so the rat could conceivably use that to modulate their trial initiation time. All other analyses (sensitivity to the previous reward and previous reward regression) were performed as described above.

## Acknowledgments

We thank Paul Glimcher, Catherine Hartley, Roozbeh Kiani, Kenway Louie, Kevin Miller, Cristina Savin and members of the Constantinople lab for feedback. We thank Madori Spiker, Daljit Kaur, Mitzi Adler-Wachter, Royall McMahon Ward, and Luke Chen for animal training.

## Author Contributions

C.M.C. designed the task. V.B. wrote training software. V.B. and C.M.C. developed software infrastructure for the high-throughput facility and data management. A.M. and C.M.C. analyzed the data. S.S.S. contributed to behavioral experiments. A.M. prepared the figures. C.M.C. and A.M. wrote the manuscript. C.M.C supervised the project.

## Supplementary materials

**Extended Data Fig. 1:**
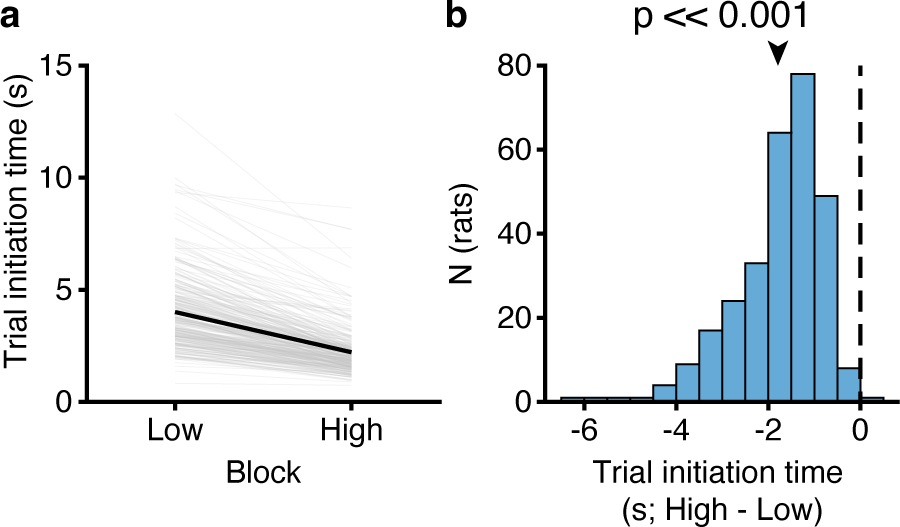
Trial initiation time in units of seconds. **a.** Trial initiation time by block (*N* = 291). Data are replotted from Fig. 1h but in units of seconds. **b.** Trial initiation time difference (high - low) across all rats. (Wilcoxon signed-rank test, *N* = 291).

**Extended Data Fig. 2:**
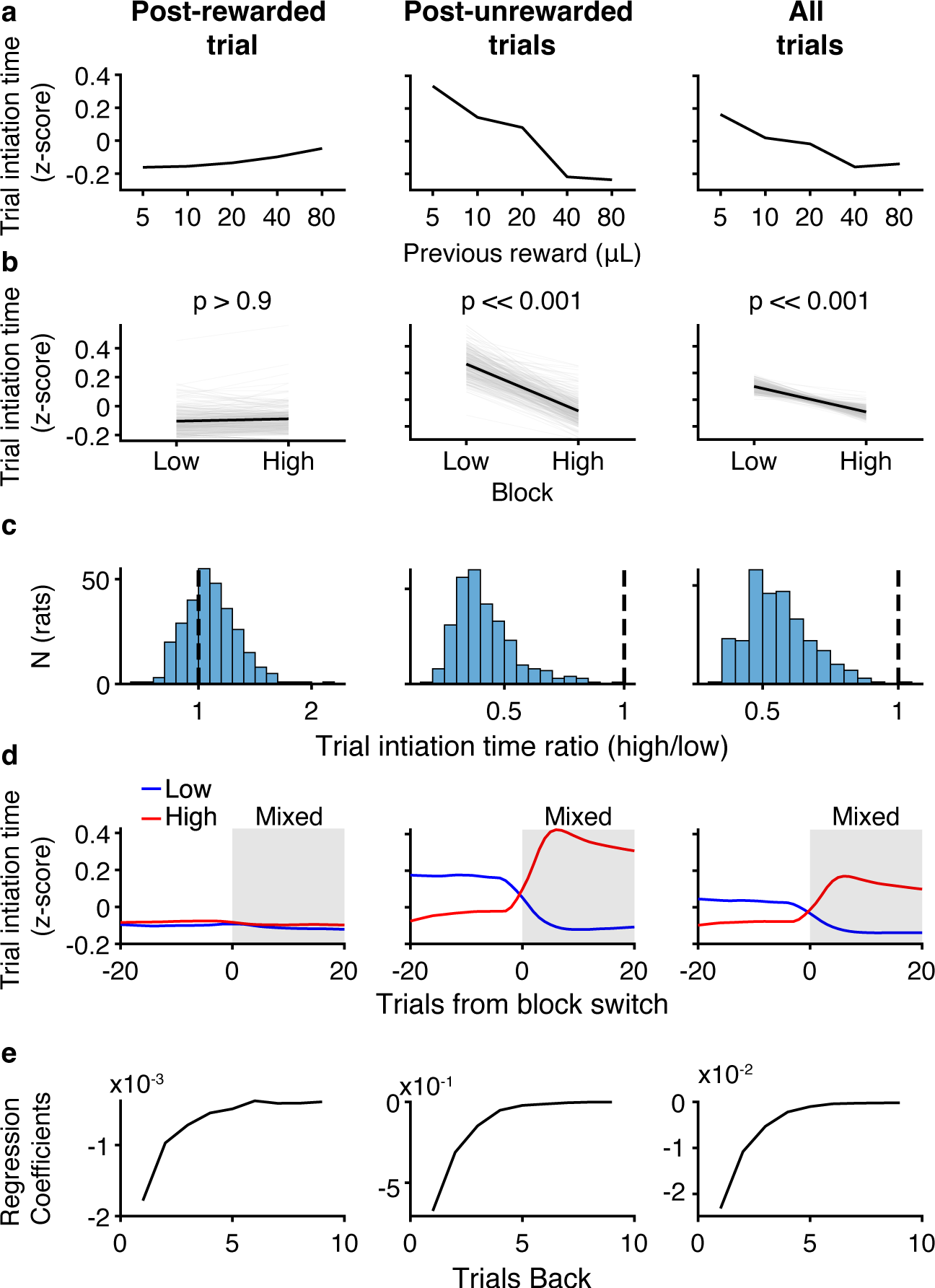
Trial initiation times depend on previous trial outcome. **a.** Trial initiation time by previous reward in mixed blocks for (left) post-rewarded trials, (center) post-unrewarded trials, and (right) all trial. **b.** Trial initiation time averaged over block (Wilcoxon Signed-rank test, N = 291). **c.** Trial initiation time ratio (mean trial initiation time in high blocks/low blocks, *N* = 291). **d.** Mean change trial initiation times from low or high blocks to mixed blocks, *N* = 291. **e.** Previous trial regression coefficients in mixed blocks, *N* = 291.

**Extended Data Fig. 3:**
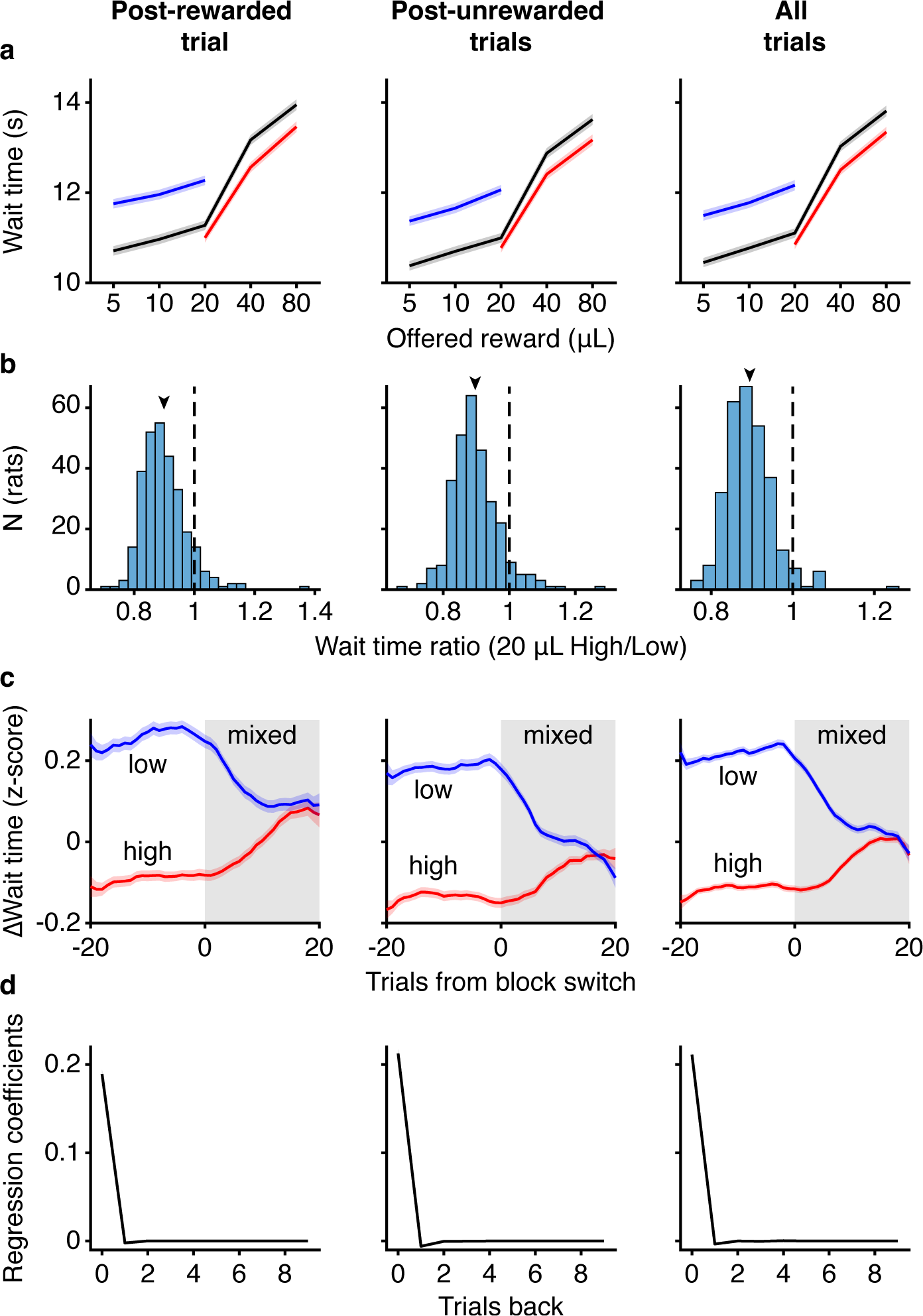
Wait times are not affected by previous trial outcome. **a.** Average wait time by volume for each block conditioned on whether the previous trial was (left) rewarded or (center) unrewarded, and (right) all trials (*N* = 291). **b.** Wait time ratios (wait time for 20 *µ*L High/Low) across rats (*N* = 291). **c.** Wait time dynamics transitioning from low (blue) or high (red) blocks into mixed blocks (*N* = 291). **d.** Reward

**Extended Data Fig. 4:**
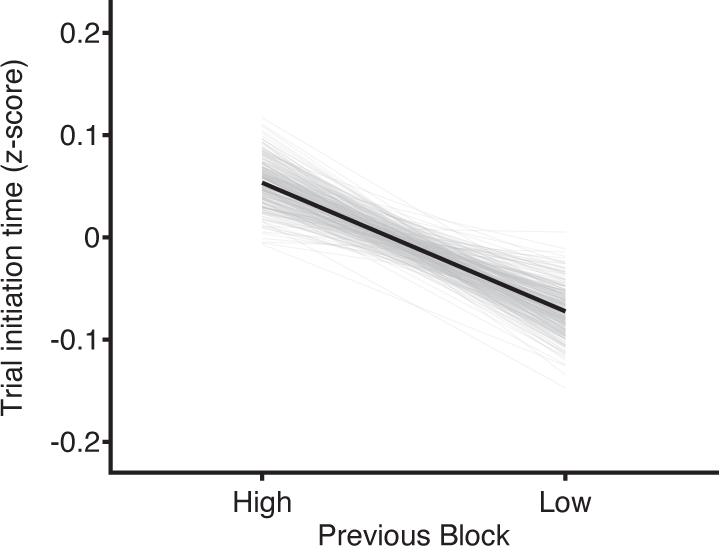
Average trial initiation time in mixed blocks conditioned on the previous block. (*N* = 291).

**Extended Data Fig. 5:**
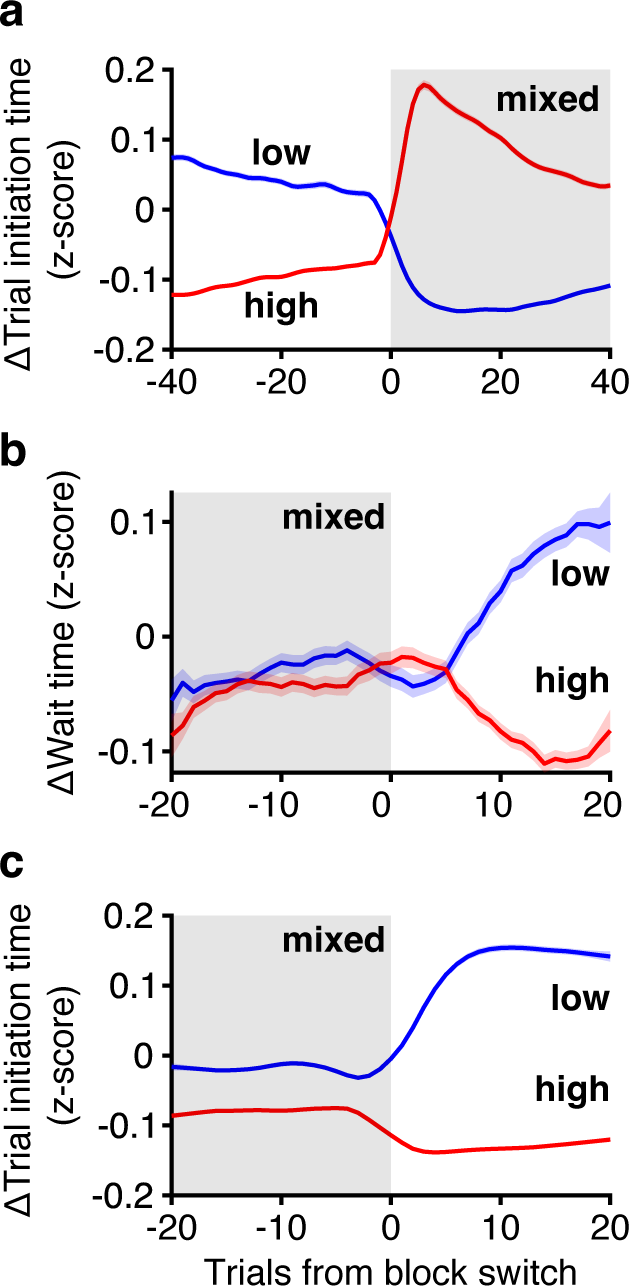
Dynamics of wait times (top) and trial initiation times (bottom) at transitions from mixed to high (red) or low (blue) blocks. **a.** Data are replotted from Fig. 2b, but with expanded x-axis limits. Trial initiation times still maintain contrast effects 40 trials into mixed blocks. **b.** Wait time transitions from mixed to high (red) and low (blue) blocks. **c.** Trial initiation time transitions from mixed to high (red) and low (blue). Block labels refer to the block at trial 0 after the mixed block. Colors are flipped relative to Fig. 2b because a current low block (blue here) is always preceded by a high block (red in Fig. 2b).

**Extended Data Fig. 6:**
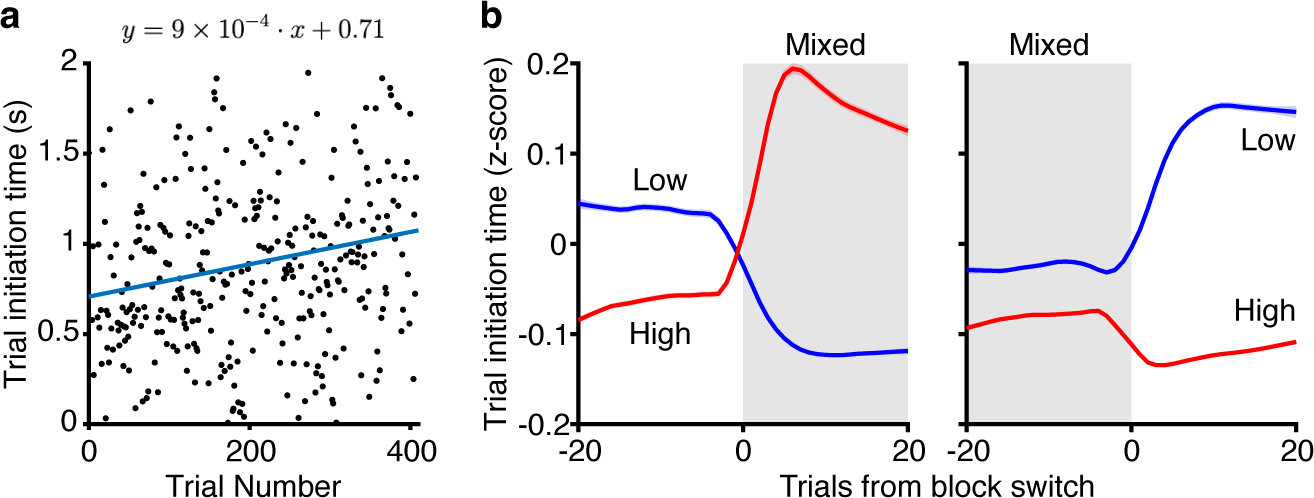
Satiety effects for trial initiation time are modest and do not qualita-tively affect results. **a.** Trial initiation time for an example session as a function of trial number. Line is least-squares regression. **b.** Trial initiation times block transition plots without detrending. Results are qualitatively similar to Fig. 2.

**Extended Data Fig. 7:**
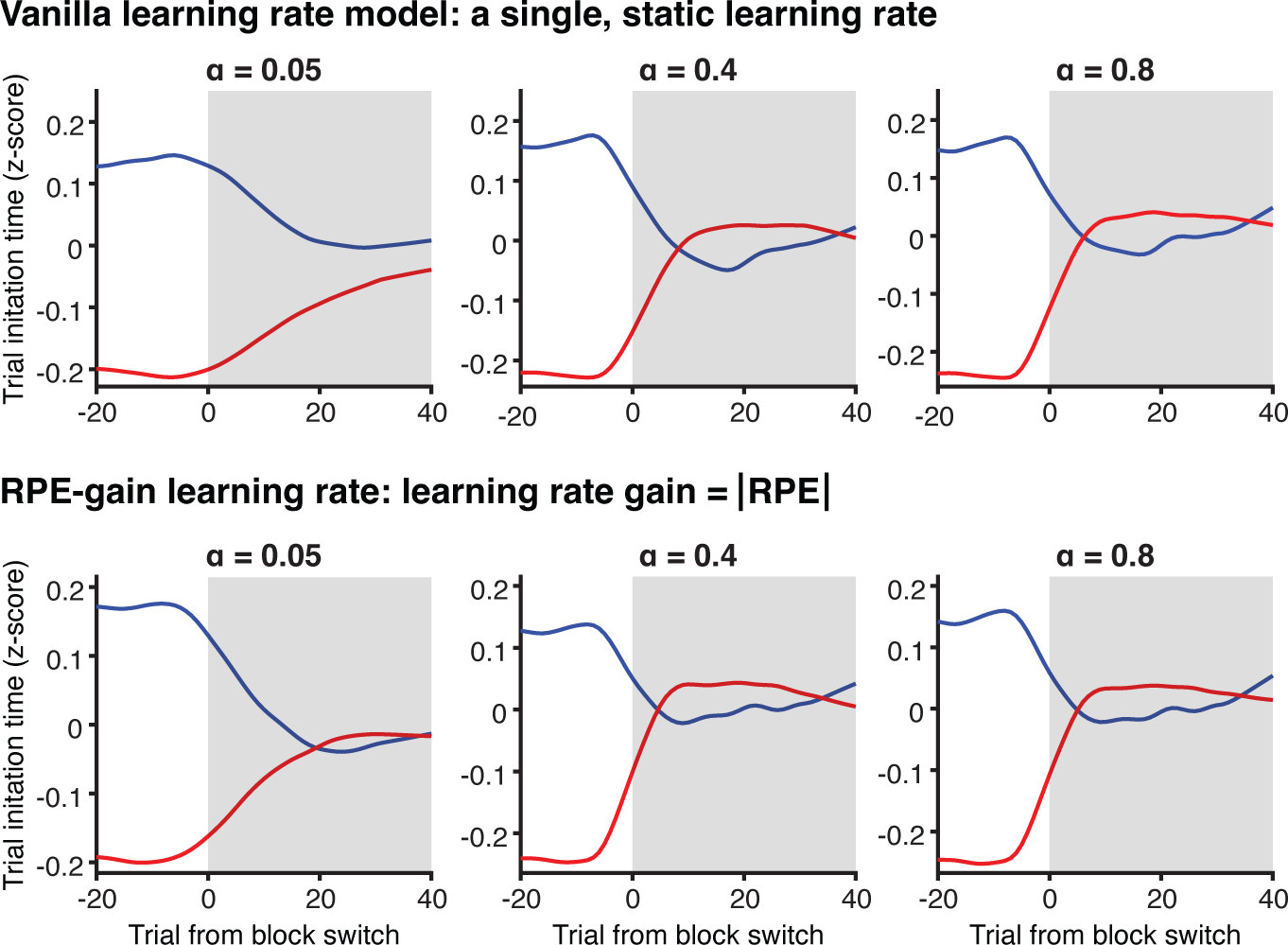
Alternative retrospective models fail to capture both fast and slow trial initiation time dynamics at block transitions. Trial initiation time model transitions from low (blue) or high (red) blocks to mixed blocks. Top: A “vanilla” learning rate model with a single, static learning rate. Bottom: a dynamic learning rate model where learning rate gain is equal to the unsigned RPE of that trial.

**Extended Data Fig. 8:**
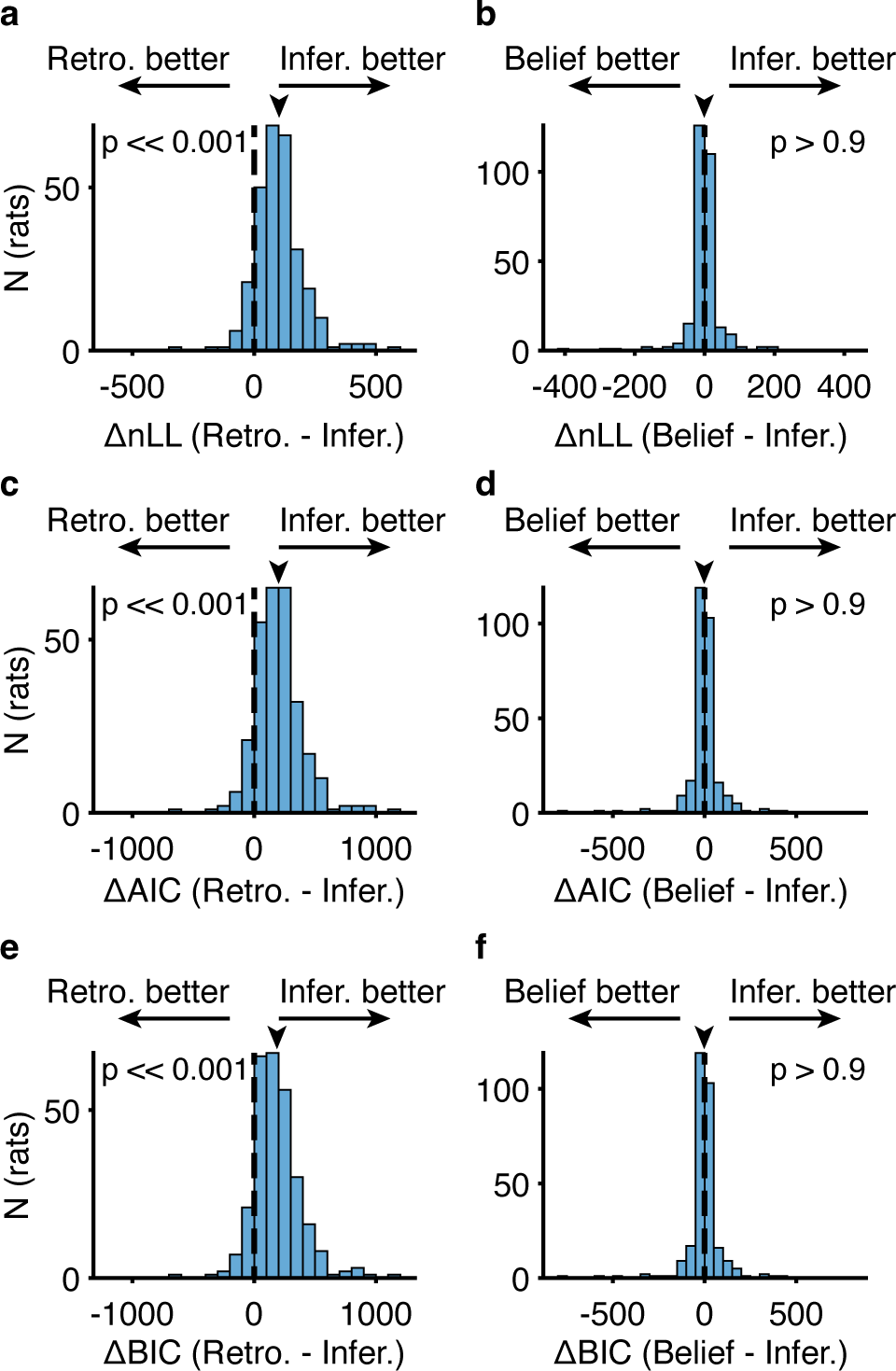
Model comparison for wait times favors inferential over retrospective model, but does not distinguish between inferential and belief state models. **a-b.** Cross-validated negative log-likelihood comparing inferential model and (a.) retrospective or (b.) belief state model. **c-d.** Akaike information criterion (AIC) comparing inferential model and (c.) retrospective or (d) belief state model. **e-f.** Bayesian information criterion (BIC) comparing inferential model and (e.) retrospective or (f.) belief state model. For each, Wilcoxon signed-rank test, *N* = 291

**Extended Data Fig. 9:**
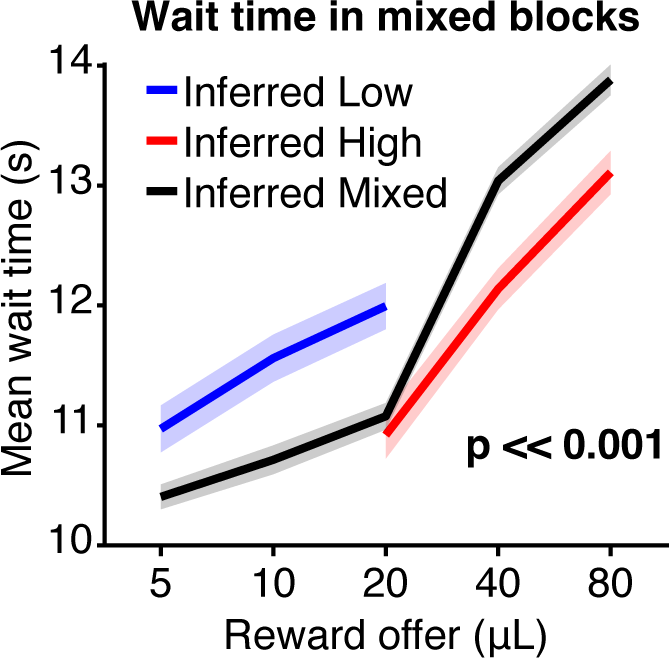
Inferential model identifies mistaken inferences during mixed blocks across rats. **a.** Average wait time curves conditioned by model-inferred block in mixed blocks only in held-out test set across rats. **b.** Wait time ratio (wait time on 20 *µ*L inferred high/low trials) is modulated by inferred block (*p <<* 0.001, Wilcoxon Signed-rank test, *N* = 291)

**Extended Data Fig. 10:**
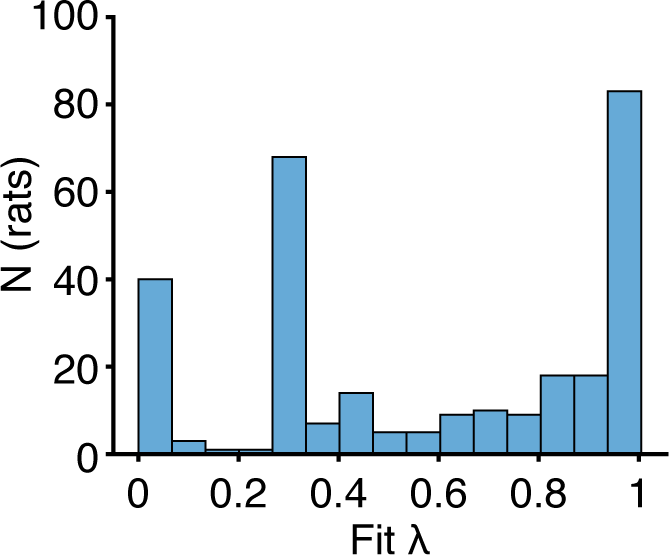
Sub-optimal inferential model with lambda. Distribution of *λ* fit over rats (*N* = 291).

**Extended Data Fig. 11:**
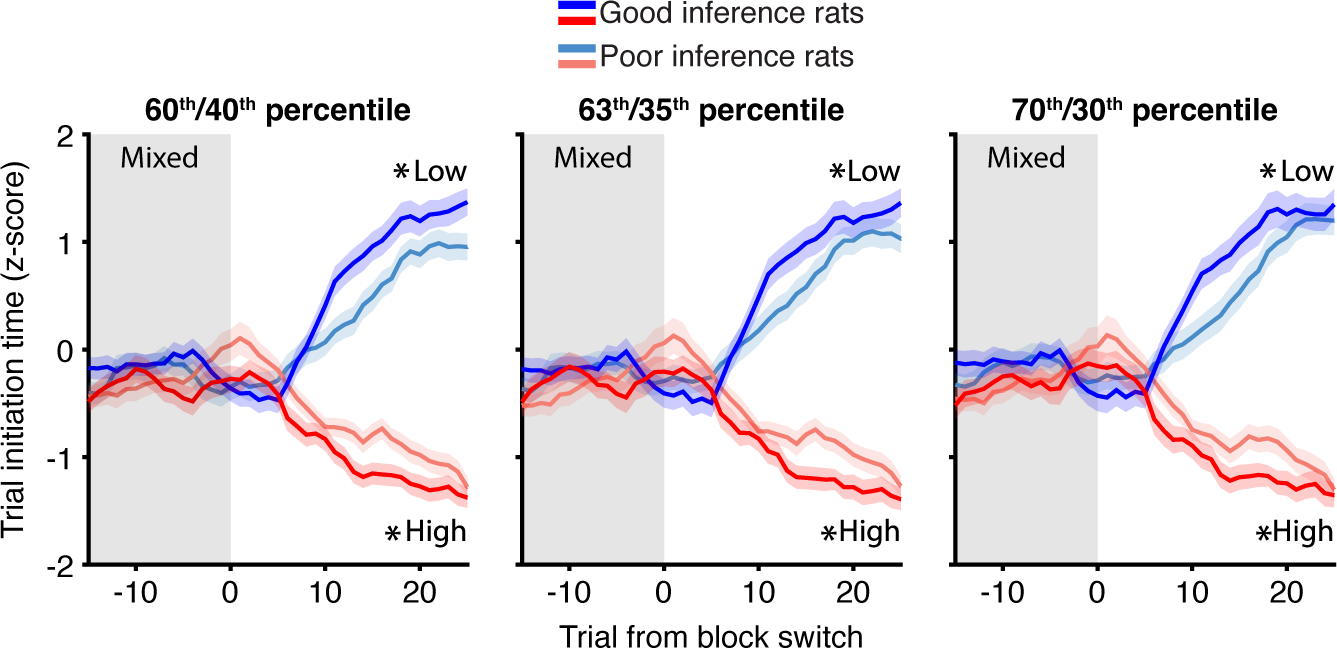
Differential wait time dynamics based on *λ* from sub-optimal Bayes model are robust across a range of percentiles.

**Extended Data Fig. 12:**
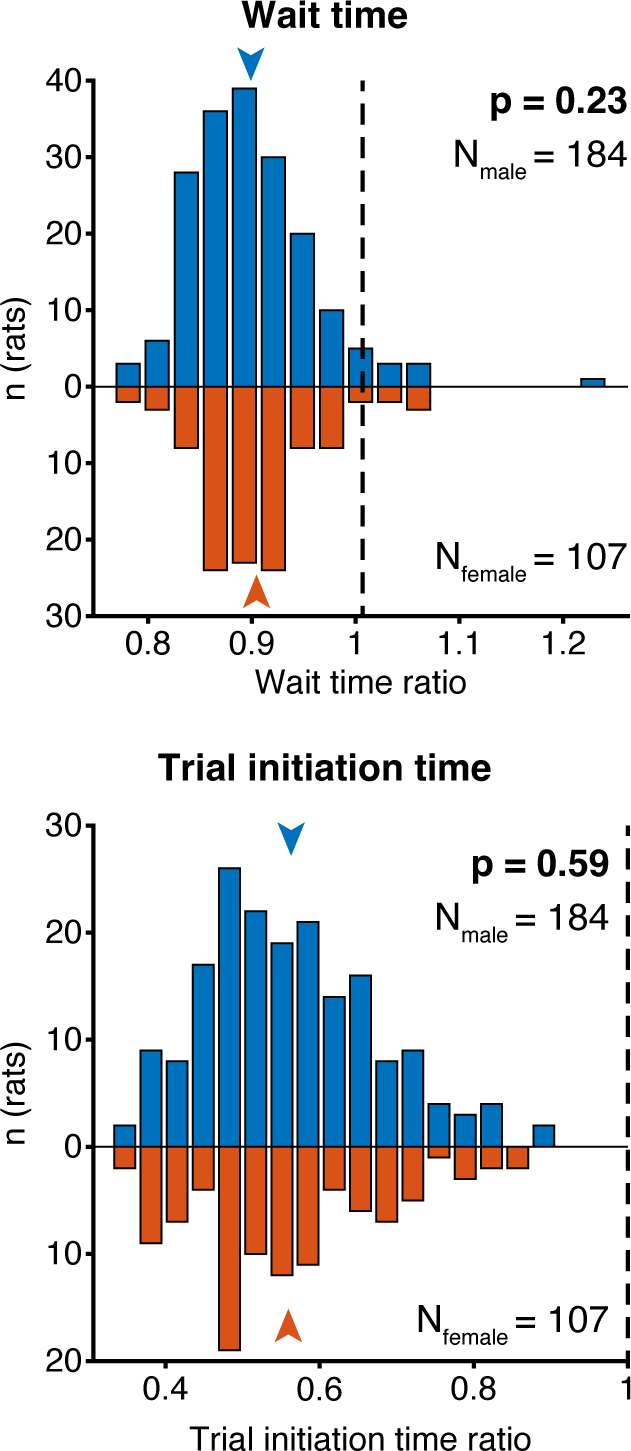
Males and females have comparable wait time ratios (top) and trial initiation time ratios (bottom) Wait time *p* = 0.23, Wilcoxon Rank-sum test, N = 184 males, 107 females. Trial initiation time *p* = 0.59, Wilcoxon Rank-sum test, N = 184 males, 107 females.

**Extended Data Fig. 13:**
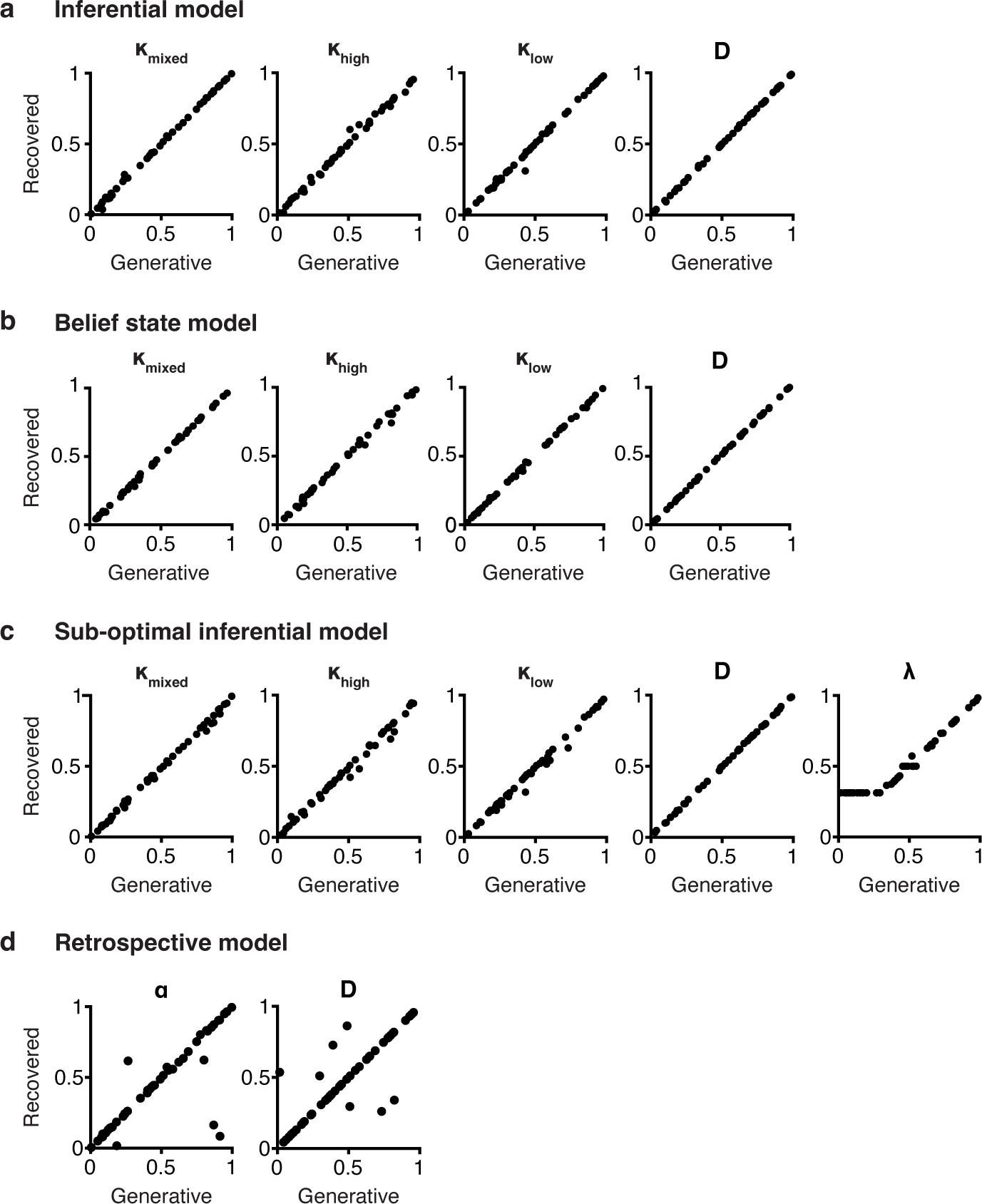
Models are able to recover generative parameters. *N* = 48 random parameter sets.

**Extended Data Fig. 14:**
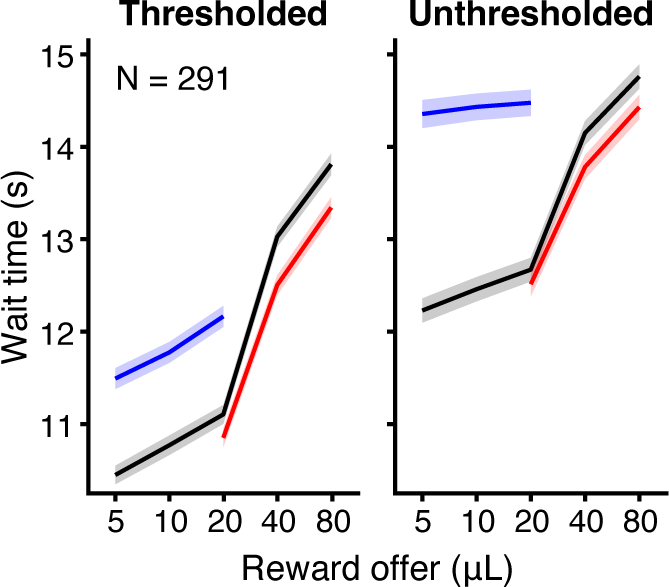
Wait time curves without threshold (right) have qualitatively similar context effects, but longer average wait times. Wait times one standard deviation above the pooled session mean were excluded for most analyses in this study (left). Including all wait times preserved the contextual effects, but resulted in longer average wait times, as the mean is particularly sensitive to outliers. Outlier wait times tended to occur in low blocks, likely due to attentional or motivational lapses. Therefore, the main difference between the thresholded and unthresholded data is that the wait time curves in low blocks are both flatter and longer in the unthresholded data.

